# Estimating spatially variable and density-dependent survival using open-population spatial capture-recapture models

**DOI:** 10.1101/2022.03.04.482982

**Authors:** Cyril Milleret, Soumen Dey, Pierre Dupont, Daniel Turek, Perry de Valpine, Richard Bischof

## Abstract

Open-population spatial capture-recapture (OPSCR) models use the spatial information contained in individual detections collected over multiple consecutive occasions to estimate occasion-specific density, but also demographic parameters. OPSCR models can also estimate spatial variation in vital rates, but such models are neither widely used nor thoroughly tested. We developed a Bayesian OSPCR model that not only accounts for spatial variation in survival using spatial covariates, but also estimates local density-dependent effects on survival within a unified framework. Using simulations, we show that OPSCR models provide sound inferences on the effect of spatial covariates on survival, including multiple competing sources of mortality, each with potentially different spatial determinants. Estimation of local density-dependent survival was possible but required more data due to the greater complexity of the model. Not accounting for spatial heterogeneity in survival led to positive bias in abundance estimates (up to 10% relative bias). We provide a set of features in R package nimbleSCR that allow computationally efficient fitting of Bayesian OPSCR models with spatially varying survival. The ability to make population-level inferences of spatial variation in survival is an essential step towards a fully spatially-explicit OPSCR model that can disentangle the role of multiple spatial drivers on population dynamics.

**Open Research statement:** code to reproduce the analysis is available on github; https://github.com/Cyril-Milleret/Public/tree/master/SpatialSurvivalOPSCR

## 1. Introduction

Spatial capture recapture (SCR) models are hierarchical models that explicitly use the spatial information contained in repeated individual detections to account for imperfect detection and estimate density (Efford 2004, Borchers and Efford 2008, Royle et al. 2014). Because SCR models are spatially explicit and accommodate various types of data (e.g., physical capture, photographic, non-invasive genetic, acoustic), they are now routinely used to analyze wildlife monitoring data. When data are collected over several consecutive occasions, open-population SCR (OPSCR) models can be used to estimate demographic rates and movement of individuals between occasions in addition to densities (Bischof et al. 2020a). Modelling individual movement between occasions can help distinguish between the different causes of individual disappearances from the population (Gardner et al. 2018). These properties make OPSCR models well-suited for drawing population-level inferences about the drivers of demographic processes.

Demographic rates, such as survival, are known to vary in time (Gaillard et al. 2000), with individual attributes (de Valpine et al. 2014), and across space (DeCesare et al. 2014). Temporal and individual variation of demographic parameters can be readily integrated in OPSCR models (Augustine et al. 2019, Bischof et al. 2020a), and the possibility of inferring spatial heterogeneity in survival using OPSCR models has been suggested (Royle et al. 2014) and applied (Chandler et al. 2018). However, the performance of models that estimate spatially-variable survival has not been thoroughly tested and their potential remains under-exploited. Estimation of spatially varying vital rates is a key step in the development of OPSCR models (Royle et al. 2014) as it will lead to a better understanding of the processes driving the spatial distribution of individuals (Pulliam 1988).

At their core, OPSCR models account for imperfect detection by using an observation process which assumes that an individual’s probability of detection is a function of distance from its activity center (AC) (Borchers and Efford 2008, Royle et al. 2014). The location of individual ACs is a latent quantity and is a representation of the center of the individual’s home range. AC locations are a key quantity of OPSCR models as they allow the estimation of density and inter-annual movement. AC locations also provide the spatial information necessary to characterize the environment in which individuals are located, and therefore its influence on survival (Chandler et al. 2018).

Density itself can be a key driver of survival (Gaillard et al. 2000). The study of density-dependent survival has often been limited to estimating the average population response to variation in overall population size through time (Bonenfant et al. 2009). However, variation in density is a spatiotemporal process and individuals within the population may not experience the same density. OPSCR models, by estimating spatio-temporal variation in density, offer a unique opportunity to study density-dependence in survival at the local scale while accounting for variation and uncertainty in both density and survival within a unified framework.

Here, we present a Bayesian OPSCR model that accounts for spatial variation in survival as a function of spatial covariates (e.g., characteristics of the landscape, resources availability) and density. We model survival using a hazard rate formulation to allow inferences on spatial variation in competing risks of mortality (Ergon et al. 2018). We quantify model performance by simulating OPSCR datasets under a wide range of scenarios. In addition, we quantify the consequence of ignoring spatial heterogeneity in mortality for OPSCR inferences. All functionalities are made available in R package nimbleSCR which provides tools for fitting efficient Bayesian (Markov chain Monte Carlo) MCMC models (de Valpine et al. 2017, NIMBLE Development Team 2019, Bischof et al. 2020b, Turek et al. 2021).

## 2. Methods

### 2.1. OPSCR model

To estimate spatial variation in mortality from live encounter and dead recoveries collected over several consecutive occasions (hereafter “years”), we built a Bayesian hierarchical state-space OPSCR model. The model is composed of four sub-models for 1) density and inter-annual movement, 2) demography, 3) live detections, and 4) dead recoveries. (Royle et al. 2014, Bischof et al. 2020a, Milleret et al. 2020, 2021, Dupont et al. 2021). We created two versions of the model. The first version can distinguish between two spatially-variable and competing causes of mortality. For example, it is possible to distinguish between culling and other causes of mortality in the case that all individuals culled are recovered dead (Bischof et al. 2020a). The second model version only considered one cause of mortality. This reflects a realistic limitation because dead recoveries are not always available in OPSCR datasets and distinguishing between multiple causes of death may not be possible.

#### 2.1.1. Spatial distribution and movement submodel

In SCR models, the location of individuals is represented by their activity centers (ACs) within the spatial domain (S). We used a Bernoulli point process with spatial intensity *λ*(***s***) to model the distribution of ACs, where ***s*** is a vector of spatial coordinates of ACs (Zhang et al. 2020). For t>1, the probability density of ***s***_*i,t*_ is conditional on the Euclidean distance to ***s***_*i,t*−1_:

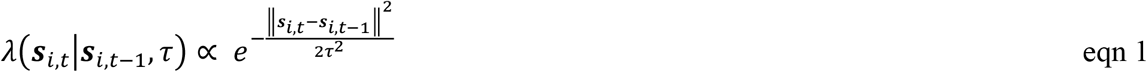

where τ is the standard deviation of a bivariate normal distribution centered on ***s***_*i,t*−1_. This represents movement between *t-1* and *t* and helps distinguish between mortality and emigration (Gardner et al. 2018).

#### 2.1.2. Demographic submodel

We used a multistate formulation (Lebreton and Pradel 2002) where each individual life history is represented by a succession of up to four discrete states z_i,t_: 1) “unborn” if the individual has not been recruited in the population; 2) “alive” if it is alive; 3) “culled” if it was culled and therefore recovered dead between the start of the previous and current occasion; or 4) “dead”: if it has died but was not recovered dead. We used data augmentation, whereby additional, undetected individuals are available for inclusion in the population at each time step (Royle et al. 2007, Royle and Dorazio 2012).

During the first year, individuals can only be designated as “unborn” (z_i,1_=1) or “alive” (z_i,1_=2) so that z_i,1_ ∼ dcat(1-*γ*_1_, *γ*_1_, 0, 0) where *γ*_1_ represents the probability to be “alive” at time *t*=1.

For *t≥2*, z_i,t_ is conditional on the state of individual *i* at *t-1:*

- If z_i,t-1_ = 1, individual *i* can be recruited (i.e., transition to state 2) with probability *γ*_t_, or remain unborn with probability 1-*γ*_t_, so z_i,t_ ∼dcat(1-*γ*_t_, *γ*_t_, 0, 0).
- If z_i,t-1_ = 2, individual *i* can survive with probability *ϕ*_*i*_ and remain z_it_=2. If it does not survive, it can either die due to culling and be recovered (transition to z_i,t_=3) with probability *h*_i_, or die from other causes without being recovered (transition to z_i,t_ =4) with probability *w*_i_, so that z_i,t_ ∼ dcat(0, *ϕ*_*i*_, h_i_, w_i_), where Φ_i_ = 1−h_i_ −w_i_.
- All individuals in dead states (z_i,t-1_ = 3 or 4) transition to z_i,t-1_ = 4, the absorbing state, with probability 1, so that z_i,t_ ∼ dcat(0, 0, 0, 1)

Abundance estimates are obtained by 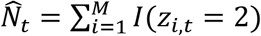, where *I*(*z*_*i,t*_ = 2) is an indicator function to count alive individuals, and *M* is the number of detected and augmented individuals.

##### Parameterization with mortality hazard rates

When an individual dies from a specific cause, it is no longer available to die from another cause. Therefore, mortality causes are competing and non-independent (Ergon et al. 2018). Following the recommendation of (Ergon et al. 2018), we parameterized the model using mortality hazard rates instead of mortality probabilities. We expressed survival and mortality probabilities as functions of the culling hazard rate (*m*_*h*_) and the hazard rate associated with to all other causes of mortality combined (*m*_*w*_). For simplicity, we assumed that the hazard rate from culling and other causes remained proportional within time intervals:

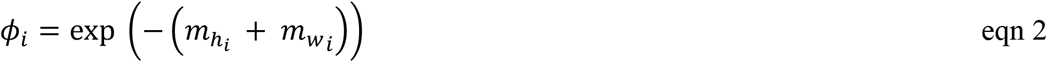

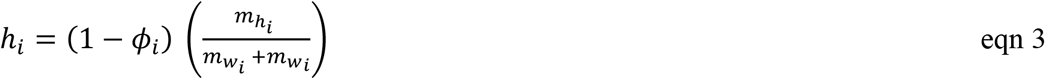

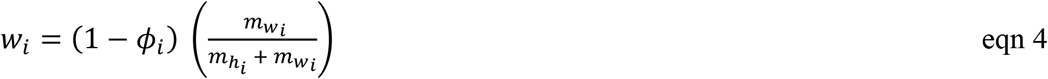

##### Spatial and individual variation in mortality

We accounted for spatial variation in cause-specific mortality by modelling mortality hazard rates (m_H_ and m_W_) as functions of a spatial covariate 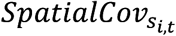 at the location of the AC (***s***_*i,t*_):

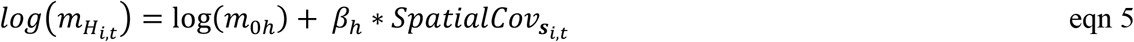

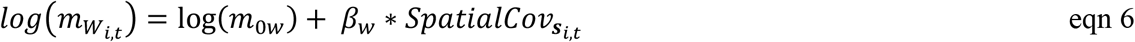

Where *β*_*h*_ and *β*_*w*_ are the coefficients of the relationship between the spatial covariate on culling and other mortality, respectively. *m*_0*h*_ and *m*_0*w*_ represent the intercept for culling and other mortality hazard rate, respectively.

##### Density dependent survival

At each occasion, local density within any habitat cell *r* of *S* (*r* =1,.., *R)* can be obtained as 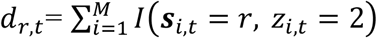, where *I*(***s***_*i,t*_ = *r, z*_*i,t*_ = 2) is an indicator function denoting whether the individual AC falls within cell *r* and it is alive.

By replacing the *SpatialCov* in Eqn.5-6 with the logarithm of the local density 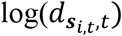, we can estimate the effect of local density-dependence on individual mortality between occasion (t-1) and t. Note that since (Eqn 5-6) are on the log scale, transformation of *d*_*r,t*_, such as *d*_*r,t*_′ = *f*(*d*_*r,t*_) = *d*_*r,t*_ + 1, is necessary to avoid 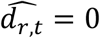.

#### 2.1.3. Live detection submodel

We used the half-normal function to model detection probability of individuals alive, whereby the probability *p*_*i,j,t*_ of detecting individual *i* at detector *j* and time *t* decreases with distance between the location x of detector j and the AC (***s***_*i,t*_):

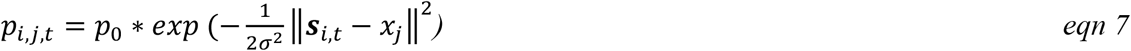

where p_0_ is the baseline detection probability, and σ the scale parameter.

The detection y_i,j,t_ of alive individual *i* at detector *j* and time *t*, is modelled as the realization of a Bernoulli process conditional on both the “alive” individual state (i.e., z_i,t_=2) and the individual and detector-specific detection probability p_i,j,t_:

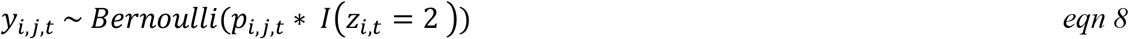

#### 2.1.4. Dead recovery model

To model dead recoveries within *S*, we used a Bernoulli point process with a bivariate normal density model, where:

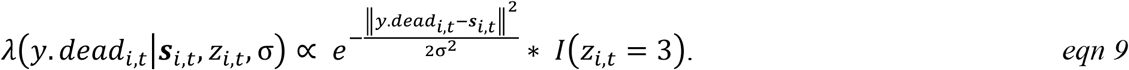

where *y*.*dead*_*i,t*_ is the vector of spatial coordinates of the dead recovery locations. The indicator function is used to condition dead recoveries on the individual being culled and recovered dead. The detection probability function represents space use, we therefore assume that σ, the shape parameter of the detection probability function, is identical for live detections and dead recoveries.

### 2.2. OPSCR model with a single cause of mortality

To provide an example of the OPSCR model where dead recoveries (*y*.*dead*) are not available and cause-specific mortality is not estimable, we built an OPSCR model with only three demographic states *z*. Individuals could be unborn (*z*_*i,t*_ = 1), alive (*z*_*i,t*_ = 2), or dead (*z*_*i,t*_ = 3). For t>1, alive individuals (*z*_*i,t*−1_ = 2) can survive with probability *ϕ*_*i,t*−1_ and remain *z*_*i,t*_ = 2 or die with probability (1 − *ϕ*_*i,t*−1_) and transition to *z*_*i,t*_ = 3, the dead absorbing state. We modelled the effect of density at occasion *t* on individual survival between occasion *t* and *t+1*:

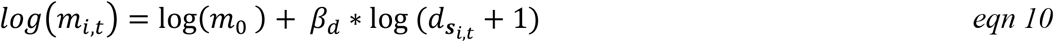

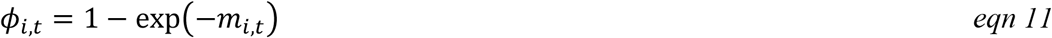

### 2.3. Simulations

We conducted simulations to quantify the performance of 1) the OPSCR model in estimating spatial variation in cause-specific mortalities with a deterministic spatial covariate; of 2) the version of the OPSCR model with integrated density-dependent survival. Finally, we tested the consequences of ignoring spatial variation in survival for abundance estimates.

#### 2.3.1. Using deterministic covariate

We created a spatial domain (S) of 28 × 28 distance units (du) subdivided in *R*=49 cells of 4×4 du. We centered in *S* a 16 × 16 detector grid (with a minimum distance of 1 du between detectors). This configuration left a 6 du buffer around the detector grid where individuals cannot be detected alive. We set *p*_0_=0.1, *σ*=2, *τ* = 3 and considered five consecutive occasions. This set-up led to an average of 2 (95% quantiles=1.5-2.3) detections per individual detected, and on average 43% (95% quantiles=34%-51%) of individuals alive detected at each occasion (Appendix 1, table 1). We created a spatially autocorrelated covariate following a diagonal gradient (*SpatialCov*, ranging from (−2 to 2); Figure 1A).

**Figure 1:**
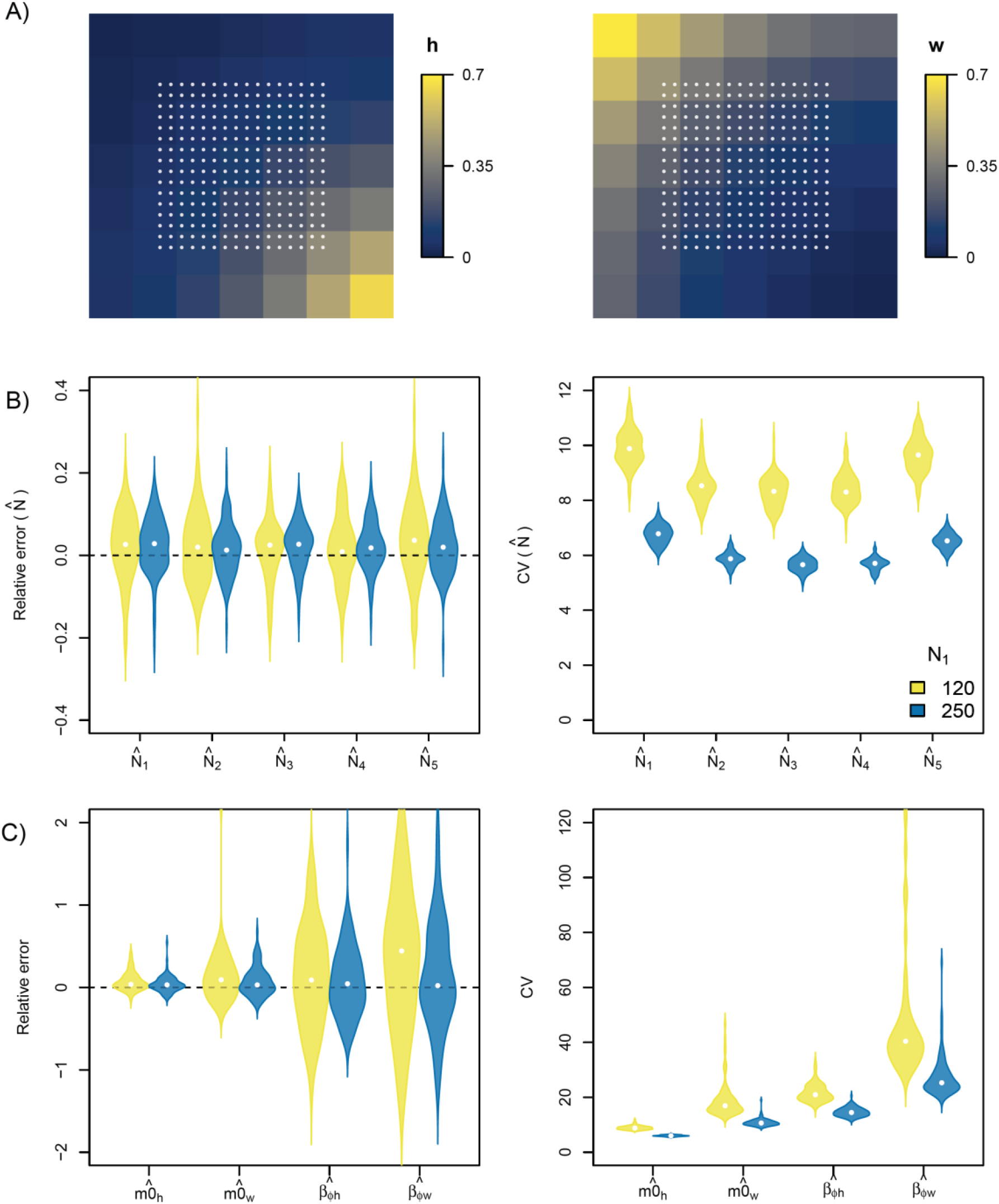
**A)** Maps depicting the simulated spatial variation in cause-specific mortality (h and w) within the spatial domain (S) for the scenario with a gradient covariate *β*_*h*_=-1 and *β*_*w*_ =1 (Table 5 Appendix 1). White points represent detectors. Violin plots show the distribution of the relative error (left; points: relative bias) and coefficient of variation (CV; right, points: median) in **B)** occasion-specific abundances, and **C)** cause-specific mortality parameters obtained after fitting the OPSCR model accounting for spatial variation in mortality to 100 replicated datasets. Results from scenarios with small (N1=120) and large (N1=250) population size are presented.

We set M=650 and N_1_= 250 individuals during the first occasion leading to 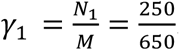. For t>1, we assumed a constant 0.3 per capita recruitment rate. We set baseline mortality hazard rates to 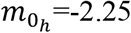 and 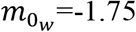 and created four different scenarios with all combinations of *β*_*h*_ = (1, −1) and *β*_*w*_ = (1, −1).

We repeated the simulation scenarios described above, but with 1) lower population size, N_1_=120 (M=370), and 2) a spatially random covariate *SpatialCov*_*r*_ ∼ *Uniform*(−1, 1) (Appendix 1, table 1). We expected estimation performance for such scenarios to be more challenging due to 1) sparser OPSCR data sets (Appendix 1, table 1), and 2) lower level of spatial autocorrelation and overall variation in mortality. In total, we simulated 100 replicated OPSCR data sets from each of the 16 scenarios (Appendix 1, table 1). We used NIMBLE’s simulation feature (de Valpine et al. 2017, NIMBLE Development Team 2019) to simulate OPSCR data sets directly from the nimble OPSCR model (Appendix 4-5).

#### 2.3.2. Using density dependent survival

To illustrate how to estimate density-dependent survival, we used the OPSCR model with a single source of mortality. Preliminary analyses showed that, computationally (convergence, mixing), OPSCR models performed more poorly when survival was modeled as dependent on latent density rather than deterministic spatial covariates. covariate. To counter this, we used a larger spatial domain (S) to increase the number of habitat cells (R) and thus provide the model with more variable latent density points to serve as covariate on the mortality hazard rate. This led to habitat of 40×40 distance units (du) subdivided in R = 64 cells of dimension 5×5 du and in which we centered a 30 × 30 detector grid. We set M=650 and N_1_ = 250. We set *m*_0_ =1.6 and *β*_*d*_ = −1 to simulate negative density-dependence on survival and simulated 100 replicated OPSCR data sets (Appendix 1, Table 12).

#### 2.3.3. Ignoring spatial heterogeneity in mortality

To evaluate whether ignoring spatial variation in mortality leads to biased parameter estimates, especially abundance 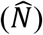, we fitted all simulated datasets described above with OPSCR models that assumed constant survival across space and time. As in the earlier simulations, we fitted models with two competing risks for the deterministic spatial covariate scenarios and a single cause of mortality for the density dependent scenario.

### 2.4. Model fitting

We fitted the Bayesian OPSCR models using Markov chain Monte Carlo (MCMC) simulation with NIMBLE in R version 4.1.0 (R Core Team 2021). We used R package nimbleSCR (Bischof et al. 2020b, Turek et al. 2021) which implements the local evaluation approach (Milleret et al. 2019) to increase MCMC efficiency. For each simulation, we ran three chains of 30,000 (60,000 for density-dependent survival) iterations, including a 2000-iteration burn-in. We considered models as converged when the Gelman-Rubin diagnostic (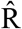, (Gelman and Rubin 1992)) was < 1.1 for all parameters and by visually inspecting trace plots from a randomly selected subset of simulations. We also computed the prior-posterior distribution overlap, and used overlap ≥35% as an indicator of weak identifiability (Gimenez et al. 2009).

### 2.5. Evaluation of model performance

We summarized the posterior 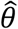 for each parameter and each simulation using relative error of the mean posterior 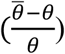 and relative precision as the coefficient of variation 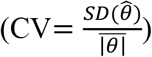, where *θ* is the true (simulated) parameter value, 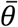 the mean of the posterior, and 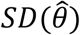 the standard deviation of the posterior. We quantified accuracy of the estimators across many simulations using relative bias of the average of the mean posteriors and the coverage accuracy of 95% credible intervals. The latter was determined as the rate of correct inferences, which is the probability that the 95% credible interval of the parameter estimate contains the true value of that parameter. In addition, we used results from a few of the simulated scenarios with a deterministic covariate (as described above) to quantify the ability of the model to predict the spatial pattern in mortality (see further details in Appendix 3).

## 3. Results

### 3.1. Spatial heterogeneity using a deterministic spatial covariate

All model parameters converged and were identifiable for scenarios with the spatial gradient covariate and large population size (N_1_=250) (Appendix 1, table 2). The combination of the spatially random covariate and low population size (N_1_=120) led to the poorest parameter convergence and identifiability (28-57%; Appendix 1, table 2).

All model parameters had low bias (≤6% relative bias) for all scenarios with large population size and a gradient covariate (Figure 1, Appendix 1, table 3-6). Accurate estimates were more challenging to obtain for scenarios with low population size and a spatially-random covariate (Appendix 1, table 3-6). For example, relative bias of *β*_*w*_ reached 44% for the scenario with low population size (N_1_=120), a spatially-random covariate and *β*_*h*_ =-1; *β*_*w*_=1; (Appendix 1, table 5). For the same scenario, but with a larger population size (N_1_=250), relative bias for *β*_*w*_ was substantially lower (RB= 2%). Coverage remained relatively high (>90%) for all scenarios (Appendix 1, table 3-6). Across all scenarios, the effect of spatial covariates on mortality cause with no dead recovery information (*β*_*w*_) was more challenging to estimate, with lower precision (approx. 2 times larger CV) than (*β*_*h*_) (Fig 1.C, Appendix 1, table 3-6).

### 3.2. Density-dependent survival

Across the 100 replicated datasets, 86 models reached convergence and showed no identifiability issues (Appendix 2, table 13). We detected a 13% positive relative bias in *ϕ*_0_ and *β*_*d*_ but coverage was >92% for all parameters (Figure 2 A, table 14).

**Figure 2:**
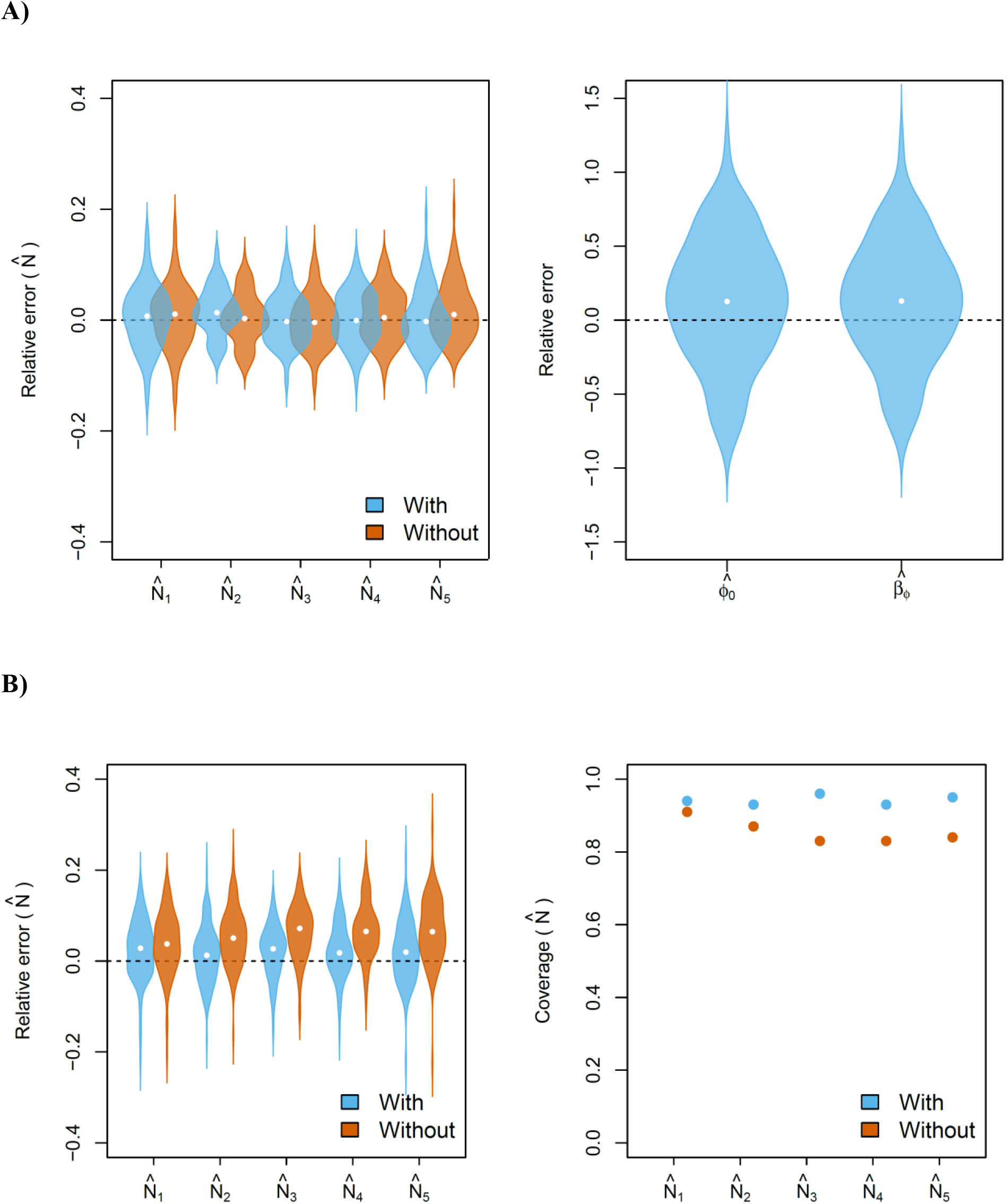
**A)** Violin plots (points: medians = relative bias) representing the distribution of the relative error in occasion-specific abundances 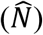 and parameters controlling for the effect of density in survival (*β*_*ϕ*_, *ϕ*_0_). Abundance estimates and associated relative error were obtained by fitting an OPSCR model that did not account for spatial variation in survival (“without”) and a model that accounted for density-dependent survival (“with”) to 100 replicated datasets simulated with a negative effect of density on survival *β*_*ϕ*_= −1. **B)** Violin plots of distribution of the relative error (left; points: medians= relative bias) and coverage (right) of occasion-specific abundances obtained by fitting an OPSCR model that accounted for spatial variation in mortality (“with) and a model did not (“without”). Results are presented for 100 replicated datasets simulated with a spatial gradient in mortality, *β*_*h*_=-1, *β*_*w*_=1 and *N*_*1*_*=250*.

### 3.3. Ignoring spatial heterogeneity in mortality

Apart from not being able to provide robust inferences on mortality, not accounting for spatial heterogeneity in mortality can lead to bias in abundance estimates. The relative bias (up to 10% for our simulated scenarios) and lower coverage were especially pronounced for scenarios with a spatial gradient in mortality (Figure 2 B, Appendix 1, table 7-11).

## 4. Discussion

We described and tested an OPSCR model that explicitly models and estimates spatial variation in survival. The model is versatile enough to allow survival to be modelled as a function of any spatial covariate, including local density estimated within the same model. Survival is modeled as a function of the location of the AC of both detected and undetected individuals which allows population-level and spatially-explicit inferences. Using simulations, we show that the model produces sound inferences on the role of spatial covariates and density dependence in explaining spatial variation in survival. In addition, the model allows for integrating spatial dead recoveries (Dupont et al. 2021) and estimates multiple competing sources of mortality with potentially different spatial determinants. The model overcomes a challenge faced by other methods, namely to a obtain population-level assessment of spatial determinants of variation in survival (Royle et al. 2018).

Despite the recognized potential of OPSCR models to estimate spatial variation in vital rates (Royle et al. 2014), few studies have attempted to use them for this purpose (Chandler et al. 2018)., Furthermore, we are not aware of any study that has evaluated the performance of OPSCR models that include spatially-explicit vital rates. Our simulations show that sound inferences in the spatial variation of cause-specific mortality can be obtained when survival is modelled as a function of a deterministic spatial covariate, even with relatively small OPSCR datasets (5 occasions, ∼50 individuals detected per occasion). OPSCR model performance (convergence, precision) was significantly higher for simulations with spatially autocorrelated survival (gradient) compared with spatially-random survival (Appendix 1, table 3-6).

One of the main advantages of SCR models is that they can estimate spatial variation in density. We showed that OPSCR models can simultaneously estimate local density and its effect on survival. This has the advantage that the uncertainty in both the number of individuals alive and their location is propagated when estimating the spatially link with survival. However, density being both a latent variable and a covariate used to explain variation in survival, it is computationally challenging (i.e., due to the increased number of model dependencies) to fit density dependent-survival OPSCR models. In Appendix 5, we showcase how centering the density covariate (*d*) can improve the mixing of MCMC chains of different model parameters.

Future research should focus on building and testing models that estimate spatial variation in other demographic parameters, i.e., recruitment, emigration and immigration, as it is an essential step to fully understand the mechanisms driving spatial heterogeneity in density and therefore population dynamics (Chandler et al. 2018). Meanwhile, it is possible to use a spatial covariate on the intensity parameter in the spatial point process submodel for the ACs to help account for spatial heterogeneity in recruitment (Zhang et al. 2020).

The OPSCR model described here can identify spatial variation in mortality and its determinants from spatial capture-recapture data, which are collected by many monitoring programs (Royle et al. 2018). Climatic conditions, resources, human activities, hunting, predation risk, and intra- and inter-specific competition are some examples of the pressures that are inherently spatial and known to impact survival and that could be studied with the model presented here. We also show that estimates of density obtained from OPSCR models can be biased (reached up to 10% in the scenarios tested) when spatial variation in survival is not accounted for. The Bayesian model written in NIMBLE and the set of features available in the nimbleSCR package will allow users to fit efficient and flexible OPSCR models. This development represents an essential step towards a fully spatially-explicit OPSCR model that can disentangle the role of spatial drivers on population dynamics (Chandler et al. 2018).

## Acknowledgements

Funded by the Norwegian Environment Agency (Miljødirektoratet), the Swedish Environmental Protection Agency (Naturvårdsverket), and the Research Council of Norway (NFR 286886). Code to reproduce the analysis are available at: https://github.com/Cyril-Milleret/Public/tree/master/SpatialSurvivalOPSCR

## Author contribution

CM, RB, PD and SD conceived and designed the study. CM implemented the analysis and wrote the manuscript with contributions from all authors. All authors gave final approval for publication.

