## Appendix 1-3 for "Estimating spatially variable and density-dependent survival using open-population spatial capture-recapture models"

### 1 Appendix 1. Spatial heterogeneity in mortality using a deterministic spatial covariate

#### 1.1 Accounting for spatial heterogeneity in mortality

Table 1: Summary of the 16 simulation scenarios explored to explain heterogeneity in spatial mortality using a spatial covariate. The average number (and standard deviation), across all 100 datasets, of individuals considered alive ($z_i = 2$), detected alive, and recovered dead ($z_i = 3$) are shown for each of the 5 consecutive occasions.

| $N_1$ | $\beta_h$ | $\beta_w$ | **Spatial Covariate** | **Alive** | | | | | **Detected** | | | | | **Dead** | | | | |
|---|---|---|---|---|---|---|---|---|---|---|---|---|---|---|---|---|---|---|
| | | | | 1 | 2 | 3 | 4 | 5 | 1 | 2 | 3 | 4 | 5 | 1 | 2 | 3 | 4 | 5 |
| 120 | 1 | 1 | Gradient | 119.6( 9.2) | 124.0( 9.2) | 125.4( 9.7) | 129.2(10.7) | 133.3( 9.5) | 47.0(6.6) | 50.5(6.5) | 52.8(7.1) | 55.6(7.3) | 57.9(7.6) | 0 | 12.3(3.3) | 12.8(3.8) | 12.7(3.2) | 12.3(3.8) |
| 120 | -1 | 1 | Gradient | 250.7(11.3) | 257.1(12.7) | 264.0(14.1) | 270.0(17.5) | 275.3(17.3) | 99.6(10.3) | 105.8( 9.7) | 112.1(10.8) | 117.7(12.2) | 120.4(11.6) | 0 | 25.7(5.4) | 25.2(5.2) | 26.6(5.0) | 26.2(5.2) |
| 120 | 1 | -1 | Gradient | 121.2(10.0) | 125.8( 9.4) | 126.3( 9.3) | 125.0( 8.8) | 127.0( 9.8) | 48.3(6.6) | 51.9(5.9) | 53.8(7.2) | 54.2(7.8) | 54.9(7.1) | 0 | 12.5(3.4) | 13.5(3.8) | 13.6(4.2) | 13.4(3.8) |
| 120 | -1 | -1 | Gradient | 248.5(10.7) | 254.3(11.4) | 260.2(13.2) | 261.4(12.8) | 263.6(13.4) | 98.0( 8.4) | 106.1( 9.7) | 111.5( 9.6) | 114.9(11.0) | 116.0(10.3) | 0 | 27.0(5.1) | 26.6(5.7) | 27.6(6.0) | 28.2(6.1) |
| 250 | 1 | 1 | Gradient | 120.0(10.2) | 123.0( 8.8) | 124.2( 8.6) | 125.9( 9.2) | 128.2(11.0) | 47.0(6.7) | 49.9(6.8) | 52.4(6.0) | 54.2(7.2) | 56.5(7.6) | 0 | 12.7(4.1) | 12.6(4.0) | 13.1(3.6) | 13.3(3.8) |
| 250 | -1 | 1 | Gradient | 248.0(10.9) | 256.5(11.1) | 263.8(13.0) | 266.9(12.3) | 266.4(14.0) | 97.3( 8.5) | 107.2( 9.0) | 111.0( 9.6) | 116.1( 8.7) | 118.2(10.8) | 0 | 26.3(5.9) | 26.3(5.8) | 28.2(5.3) | 29.0(5.6) |
| 250 | 1 | -1 | Gradient | 119.6( 8.0) | 124.5( 9.2) | 128.3( 8.7) | 131.0( 9.5) | 133.2(11.5) | 47.0(6.0) | 51.8(6.8) | 53.8(6.7) | 56.8(7.0) | 58.6(8.0) | 0 | 11.6(3.4) | 12.3(3.3) | 12.7(3.7) | 12.8(4.1) |
| 250 | -1 | -1 | Gradient | 250.2(12.3) | 258.9(12.6) | 265.6(14.5) | 271.6(17.5) | 275.2(19.2) | 99.3( 9.6) | 107.0( 9.9) | 112.0(10.4) | 117.0(12.0) | 120.5(14.3) | 0 | 25.0(5.1) | 25.9(5.4) | 26.1(5.1) | 27.1(6.0) |
| 120 | 1 | 1 | Random | 119.8(8.8) | 118.7(9.2) | 123.4(8.1) | 127.3(9.5) | 134.4(9.2) | 47.9(6.7) | 51.2(6.4) | 53.9(6.0) | 56.3(6.7) | 60.1(6.1) | 0 | 14.2(3.4) | 11.7(3.5) | 12.2(3.1) | 11.4(3.5) |
| 120 | -1 | 1 | Random | 249.1(11.5) | 248.1(11.8) | 257.5(12.5) | 268.3(13.0) | 281.6(12.6) | 99.3( 8.9) | 108.1( 9.0) | 113.6( 9.0) | 119.8( 9.9) | 126.2(11.0) | 0 | 29.1(4.6) | 24.0(5.4) | 25.4(4.8) | 24.2(5.5) |
| 120 | 1 | -1 | Random | 120.0(8.9) | 114.2(8.6) | 112.8(8.0) | 112.0(8.3) | 112.0(8.5) | 47.3(5.7) | 48.4(6.7) | 49.6(6.6) | 51.1(6.9) | 52.1(7.0) | 0 | 17.0(3.9) | 15.3(3.9) | 15.0(3.7) | 14.8(3.7) |
| 120 | -1 | -1 | Random | 251.0(13.6) | 241.5(12.2) | 237.4(12.9) | 234.4(12.3) | 232.8(10.8) | 100.1( 8.8) | 104.8( 9.4) | 107.7( 9.5) | 108.4(10.1) | 109.4( 9.4) | 0 | 34.6(5.1) | 31.3(5.1) | 31.4(5.5) | 31.5(5.5) |
| 250 | 1 | 1 | Random | 120.7(8.7) | 114.6(8.5) | 112.9(8.8) | 113.0(9.5) | 113.3(9.7) | 47.5(6.7) | 48.9(6.2) | 50.7(5.9) | 52.0(6.3) | 52.1(6.7) | 0 | 16.4(3.8) | 15.8(3.6) | 15.9(4.4) | 15.3(3.5) |
| 250 | -1 | 1 | Random | 250.5(11.6) | 239.1(11.3) | 236.9(12.5) | 236.0(12.9) | 235.5(12.3) | 99.4( 8.2) | 102.2( 8.9) | 104.1(10.3) | 105.8( 8.7) | 107.0( 9.2) | 0 | 33.2(5.2) | 31.9(5.1) | 31.5(5.8) | 32.5(6.1) |
| 250 | 1 | -1 | Random | 119.9( 8.7) | 120.5( 8.8) | 126.0(10.6) | 130.1( 9.5) | 137.1( 7.8) | 47.5(6.7) | 49.7(6.8) | 52.2(7.2) | 55.6(6.8) | 59.1(6.5) | 0 | 13.5(3.4) | 11.7(3.3) | 11.7(3.2) | 11.1(3.3) |
| 250 | -1 | -1 | Random | 249.5(13.1) | 248.9(12.4) | 261.5(11.0) | 273.7(12.0) | 285.5(10.7) | 99.5(10.2) | 104.2( 9.0) | 111.3( 9.0) | 115.4(10.0) | 120.4(10.0) | 0 | 28.7(4.7) | 23.4(4.1) | 23.7(4.5) | 24.1(4.7) |

Table 2: Percentage of models with all parameters converged, identifiable, and both converged and identifiable for the 100 datasets associated with each of the 16 scenarios tested.

| $N_1$ | $\beta_h$ | $\beta_w$ | Spatial Covariate | % Converged | % Identifiable | % Converged and Identifiable |
|---|---|---|---|---|---|---|
| 120.00 | 1.00 | 1.00 | Gradient | 100 | 100 | 100 |
| 120.00 | -1.00 | 1.00 | Gradient | 96 | 76 | 72 |
| 120.00 | 1.00 | -1.00 | Gradient | 100 | 94 | 94 |
| 120.00 | -1.00 | -1.00 | Gradient | 100 | 100 | 100 |
| 250.00 | 1.00 | 1.00 | Gradient | 100 | 100 | 100 |
| 250.00 | -1.00 | 1.00 | Gradient | 100 | 100 | 100 |
| 250.00 | 1.00 | -1.00 | Gradient | 100 | 100 | 100 |
| 250.00 | -1.00 | -1.00 | Gradient | 100 | 100 | 100 |
| 120.00 | 1.00 | 1.00 | Random | 67 | 64 | 31 |
| 120.00 | -1.00 | 1.00 | Random | 65 | 64 | 29 |
| 120.00 | 1.00 | -1.00 | Random | 78 | 79 | 57 |
| 120.00 | -1.00 | -1.00 | Random | 57 | 71 | 28 |
| 250.00 | 1.00 | 1.00 | Random | 99 | 93 | 92 |
| 250.00 | -1.00 | 1.00 | Random | 99 | 94 | 93 |
| 250.00 | 1.00 | -1.00 | Random | 98 | 92 | 90 |
| 250.00 | -1.00 | -1.00 | Random | 99 | 93 | 92 |

Table 3: Relative bias (RB) and coefficient of variation (CV) for the 100 simulated datasets for the scenario with $\beta_h$ = 1, $\beta_w$ = -1, and for a) $N_1$ = 120 and b) $N_1$ = 250. Values within brackets show the 95% quantiles.

(a) $N_1 = 120$

| | Random | | | Gradient | | |
|---|---|---|---|---|---|---|
| Parameters | RB | CV | Coverage | RB | CV | Coverage |
| | | | $\beta_h = 1$ $\beta_w$ = -1 | | | |
| $N_1$ | 0.00 (-0.18-0.22) | 10.29 (8.60-11.84) | 93 | 0.02 (-0.20-0.18) | 9.80 (8.69-11.33) | 97 |
| $N_2$ | 0.02 (-0.14-0.19) | 8.74 (7.51-10.28) | 97 | 0.03 (-0.13-0.16) | 8.56 (7.63-9.79) | 98 |
| $N_3$ | 0.02 (-0.10-0.17) | 8.42 (7.23-9.36) | 97 | 0.04 (-0.09-0.21) | 8.17 (7.24-9.04) | 93 |
| $N_4$ | 0.02 (-0.11-0.20) | 8.42 (7.56-9.63) | 91 | 0.04 (-0.08-0.15) | 8.15 (7.33-9.08) | 99 |
| $N_5$ | 0.03 (-0.17-0.18) | 9.38 (8.45-10.74) | 96 | 0.05 (-0.09-0.19) | 9.39 (8.44-10.94) | 98 |
| $\beta_h$ | 0.04 (-1.05-1.65) | 55.69 (29.71-1414.65) | 100 | -0.01 (-0.41-0.49) | 25.12 (19.90-36.51) | 100 |
| $\beta_w$ | -0.06 (-1.80-1.54) | 87.26 (46.80-913.09) | 100 | 0.13 (-0.52-0.82) | 33.68 (25.79-63.49) | 100 |
| $\tau$ | 0.00 (-0.16-0.12) | 7.24 (6.31-9.34) | 100 | -0.01 (-0.12-0.15) | 7.33 (6.28-8.72) | 100 |
| $mh_h$ | 0.07 (-0.13-0.32) | 10.51 (7.70-16.79) | 100 | 0.02 (-0.17-0.22) | 9.11 (8.15-10.61) | 100 |
| $mh_w$ | 0.04 (-0.24-0.52) | 21.29 (14.89-35.62) | 100 | 0.03 (-0.21-0.43) | 16.68 (14.25-23.74) | 100 |
| $p_0$ | 0.01 (-0.13-0.15) | 8.01 (7.34-9.14) | 100 | 0.01 (-0.14-0.16) | 8.10 (7.30-8.87) | 100 |
| $\sigma$ | -0.01 (-0.07-0.08) | 3.73 (3.39-4.38) | 100 | 0.00 (-0.08-0.07) | 3.75 (3.35-4.19) | 100 |

(b) $N_1 = 250$

| | Random | | | Gradient | | |
|---|---|---|---|---|---|---|
| Parameters | RB | CV | Coverage | RB | CV | Coverage |
| | | | $\beta_h = 1$ $\beta_w$ = -1 | | | |
| $N_1$ | 0.00 (-0.12-0.16) | 7.14 (6.46-7.68) | 94 | 0.01 (-0.10-0.14) | 6.81 (6.40-7.45) | 95 |
| $N_2$ | 0.02 (-0.08-0.16) | 6.00 (5.52-6.50) | 94 | 0.01 (-0.08-0.12) | 5.95 (5.56-6.40) | 97 |
| $N_3$ | 0.01 (-0.08-0.11) | 5.65 (5.23-6.29) | 97 | 0.01 (-0.11-0.12) | 5.69 (5.24-6.32) | 92 |
| $N_4$ | 0.03 (-0.07-0.15) | 5.66 (5.28-6.16) | 95 | 0.02 (-0.08-0.14) | 5.73 (5.21-6.12) | 93 |
| $N_5$ | 0.03 (-0.09-0.13) | 6.32 (5.91-6.96) | 97 | 0.01 (-0.10-0.14) | 6.54 (6.01-7.16) | 95 |
| $\beta_h$ | 0.02 (-0.83-1.04) | 40.95 (22.96-237.76) | 100 | 0.00 (-0.31-0.37) | 17.35 (14.73-22.84) | 100 |
| $\beta_w$ | 0.17 (-1.30-1.27) | 55.96 (31.39-528.97) | 100 | -0.02 (-0.42-0.63) | 23.56 (18.30-35.05) | 100 |
| $\tau$ | 0.01 (-0.10-0.10) | 4.92 (4.47-5.56) | 100 | -0.01 (-0.11-0.09) | 5.12 (4.54-5.72) | 100 |
| $mh_h$ | 0.04 (-0.09-0.21) | 7.08 (5.34-12.51) | 100 | 0.00 (-0.08-0.15) | 6.29 (5.69-6.86) | 100 |
| $mh_w$ | 0.08 (-0.18-0.39) | 13.66 (9.96-22.01) | 100 | 0.01 (-0.16-0.28) | 10.90 (9.58-13.09) | 100 |
| $p_0$ | 0.01 (-0.07-0.11) | 5.48 (5.19-5.88) | 100 | -0.02 (-0.10-0.14) | 5.65 (5.28-6.12) | 100 |
| $\sigma$ | -0.01 (-0.05-0.05) | 2.55 (2.39-2.74) | 100 | 0.00 (-0.06-0.05) | 2.62 (2.42-2.89) | 100 |

Table 4: Relative bias (RB) and coefficient of variation (CV) for the 100 simulated datasets for the scenario with $\beta_h$ = -1, $\beta_w$ = -1, and for a) $N_1 = 120$ and b) $N_1 = 250$. Values within brackets show the 95% quantiles.

(a) $N_1 = 120$

| | **Random** | | | **Gradient** | | |
|---|---|---|---|---|---|---|
| Parameters | RB | CV | Coverage | RB | CV | Coverage |
| $\beta_h$ = -1 $\beta_w$ = -1 | | | | | | |
| $N_1$ | 0.01 (-0.18-0.21) | 10.22 (8.98-11.82) | 96 | 0.03 (-0.17-0.20) | 9.77 (8.54-11.37) | 98 |
| $N_2$ | 0.03 (-0.19-0.17) | 8.62 (7.63-9.78) | 91 | 0.02 (-0.15-0.18) | 8.45 (7.47-9.62) | 93 |
| $N_3$ | 0.01 (-0.10-0.18) | 8.18 (7.53-9.11) | 96 | 0.03 (-0.13-0.16) | 7.98 (7.06-9.46) | 97 |
| $N_4$ | 0.02 (-0.08-0.18) | 8.29 (7.31-9.07) | 95 | 0.03 (-0.11-0.18) | 7.99 (7.01-9.30) | 93 |
| $N_5$ | 0.04 (-0.14-0.21) | 9.27 (7.96-10.23) | 96 | 0.05 (-0.16-0.20) | 8.80 (7.77-10.09) | 94 |
| $\beta_h$ | -0.03 (-1.20-1.93) | 58.55 (33.61-881.07) | 100 | 0.02 (-0.31-0.45) | 22.45 (18.22-30.19) | 100 |
| $\beta_w$ | 0.07 (-1.64-1.51) | 85.12 (46.30-735.50) | 100 | 0.15 (-0.49-1.05) | 32.91 (25.01-59.95) | 100 |
| $\tau$ | -0.02 (-0.15-0.10) | 7.34 (6.09-8.93) | 100 | 0.00 (-0.15-0.11) | 7.24 (6.20-9.25) | 100 |
| $mh_h$ | 0.05 (-0.11-0.39) | 9.18 (7.53-19.58) | 100 | 0.01 (-0.13-0.17) | 7.99 (7.27-8.93) | 100 |
| $mh_w$ | 0.08 (-0.17-0.55) | 20.50 (14.98-34.06) | 100 | 0.06 (-0.23-0.57) | 16.46 (14.00-25.56) | 100 |
| $p_0$ | 0.02 (-0.13-0.15) | 7.87 (7.14-8.79) | 100 | 0.00 (-0.14-0.14) | 7.93 (7.16-8.99) | 100 |
| $\sigma$ | -0.01 (-0.07-0.06) | 3.71 (3.35-4.15) | 100 | 0.00 (-0.06-0.07) | 3.72 (3.32-4.29) | 100 |

(b) $N_1 = 250$

| | **Random** | | | **Gradient** | | |
|---|---|---|---|---|---|---|
| Parameters | RB | CV | Coverage | RB | CV | Coverage |
| $\beta_h$ = -1 $\beta_w$ = -1 | | | | | | |
| $N_1$ | 0.02 (-0.13-0.15) | 7.07 (6.49-7.87) | 94 | 0.03 (-0.10-0.17) | 6.95 (6.20-7.61) | 91 |
| $N_2$ | 0.01 (-0.11-0.14) | 5.95 (5.40-6.59) | 95 | 0.02 (-0.09-0.14) | 5.87 (5.40-6.37) | 93 |
| $N_3$ | 0.01 (-0.10-0.10) | 5.64 (5.32-6.31) | 97 | 0.03 (-0.08-0.15) | 5.54 (5.07-5.92) | 91 |
| $N_4$ | 0.01 (-0.10-0.12) | 5.62 (5.09-6.29) | 97 | 0.02 (-0.10-0.13) | 5.52 (5.04-6.10) | 90 |
| $N_5$ | 0.01 (-0.12-0.15) | 6.26 (5.65-7.10) | 92 | 0.01 (-0.09-0.12) | 6.13 (5.51-6.78) | 96 |
| $\beta_h$ | 0.13 (-0.73-1.07) | 36.93 (22.78-138.11) | 100 | 0.03 (-0.28-0.33) | 15.49 (13.14-20.25) | 100 |
| $\beta_w$ | 0.11 (-1.09-1.57) | 54.58 (32.03-612.74) | 100 | 0.03 (-0.40-0.72) | 22.55 (17.58-32.78) | 100 |
| $\tau$ | 0.00 (-0.10-0.09) | 5.01 (4.41-5.64) | 100 | 0.00 (-0.09-0.10) | 4.96 (4.43-5.63) | 100 |
| $mh_h$ | 0.02 (-0.08-0.19) | 6.31 (5.12-10.94) | 100 | 0.01 (-0.10-0.12) | 5.51 (5.11-5.97) | 100 |
| $mh_w$ | 0.07 (-0.21-0.42) | 13.23 (9.75-25.22) | 100 | 0.02 (-0.20-0.35) | 10.95 (9.82-13.29) | 100 |
| $p_0$ | 0.00 (-0.09-0.12) | 5.46 (5.02-5.91) | 100 | -0.01 (-0.12-0.07) | 5.53 (5.20-6.07) | 100 |
| $\sigma$ | 0.00 (-0.04-0.05) | 2.57 (2.33-2.83) | 100 | 0.00 (-0.05-0.05) | 2.58 (2.40-2.84) | 100 |

Table 5: Relative bias (RB) and coefficient of variation (CV) for the 100 simulated datasets for the scenario with $\beta_h$ = -1, $\beta_w$ = 1, and for a) $N_1 = 120$ and b) $N_1 = 250$. Values within brackets show the 95% quantiles.

(a) $N_1 = 120$

| | **Random** | | | **Gradient** | | |
|---|---|---|---|---|---|---|
| Parameters | RB | CV | Coverage | RB | CV | Coverage |
| $\beta_h$ = -1 $\beta_w = 1$ | | | | | | |
| $N_1$ | 0.04 (-0.12-0.16) | 9.91 (8.80-11.69) | 100 | 0.03 (-0.18-0.14) | 9.88 (8.79-11.28) | 95 |
| $N_2$ | 0.03 (-0.10-0.17) | 8.41 (7.50-9.39) | 96 | 0.02 (-0.11-0.21) | 8.53 (7.55-9.94) | 95 |
| $N_3$ | 0.05 (-0.18-0.17) | 8.15 (7.12-9.52) | 93 | 0.02 (-0.16-0.15) | 8.33 (7.41-9.16) | 95 |
| $N_4$ | 0.02 (-0.13-0.19) | 8.25 (7.35-9.45) | 95 | 0.01 (-0.12-0.17) | 8.30 (7.64-9.71) | 98 |
| $N_5$ | 0.02 (-0.19-0.20) | 9.36 (8.54-10.65) | 93 | 0.04 (-0.17-0.21) | 9.65 (8.39-10.79) | 96 |
| $\beta_h$ | 0.09 (-0.82-1.35) | 57.05 (32.58-271.78) | 100 | 0.01 (-0.37-0.60) | 20.94 (17.58-30.28) | 100 |
| $\beta_w$ | 0.44 (-1.20-2.26) | 69.30 (37.07-440.05) | 100 | 0.15 (-0.72-1.72) | 40.41 (29.57-122.99) | 100 |
| $\tau$ | -0.01 (-0.15-0.15) | 7.18 (6.36-8.31) | 100 | 0.00 (-0.15-0.13) | 7.39 (6.47-8.49) | 100 |
| $mh_h$ | 0.04 (-0.07-0.34) | 10.80 (7.71-18.84) | 100 | 0.04 (-0.15-0.23) | 8.80 (7.89-10.88) | 100 |
| $mh_w$ | 0.09 (-0.22-0.61) | 22.13 (15.16-37.25) | 100 | 0.03 (-0.25-0.66) | 16.92 (13.76-38.40) | 100 |
| $p_0$ | -0.01 (-0.15-0.13) | 7.93 (7.23-8.67) | 100 | 0.00 (-0.16-0.20) | 8.07 (7.34-9.05) | 100 |
| $\sigma$ | 0.01 (-0.06-0.07) | 3.75 (3.33-4.17) | 100 | -0.01 (-0.08-0.08) | 3.77 (3.41-4.20) | 100 |

(b) $N_1 = 250$

| | **Random** | | | **Gradient** | | |
|---|---|---|---|---|---|---|
| Parameters | RB | CV | Coverage | RB | CV | Coverage |
| $\beta_h$ = -1 $\beta_w = 1$ | | | | | | |
| $N_1$ | 0.02 (-0.10-0.13) | 7.03 (6.56-7.79) | 98 | 0.03 (-0.13-0.14) | 6.78 (6.16-7.39) | 94 |
| $N_2$ | 0.03 (-0.13-0.14) | 6.01 (5.57-6.62) | 90 | 0.01 (-0.09-0.13) | 5.87 (5.39-6.40) | 93 |
| $N_3$ | 0.03 (-0.10-0.13) | 5.70 (5.18-6.29) | 96 | 0.03 (-0.09-0.12) | 5.66 (5.17-6.10) | 96 |
| $N_4$ | 0.03 (-0.09-0.14) | 5.70 (5.20-6.32) | 91 | 0.02 (-0.10-0.13) | 5.71 (5.20-6.20) | 93 |
| $N_5$ | 0.02 (-0.12-0.15) | 6.43 (5.91-7.01) | 90 | 0.02 (-0.08-0.14) | 6.53 (5.98-7.07) | 95 |
| $\beta_h$ | 0.04 (-0.57-0.82) | 39.67 (24.41-96.64) | 100 | -0.02 (-0.23-0.37) | 14.47 (12.23-17.78) | 100 |
| $\beta_w$ | 0.02 (-1.01-1.62) | 52.19 (31.31-471.12) | 100 | 0.08 (-0.46-0.71) | 25.28 (20.93-46.17) | 100 |
| $\tau$ | 0.01 (-0.10-0.11) | 5.02 (4.50-5.74) | 100 | -0.01 (-0.10-0.09) | 4.99 (4.47-5.69) | 100 |
| $mh_h$ | 0.03 (-0.11-0.23) | 7.14 (5.45-11.84) | 100 | 0.02 (-0.12-0.16) | 5.99 (5.49-6.52) | 100 |
| $mh_w$ | 0.03 (-0.18-0.48) | 14.53 (10.09-25.79) | 100 | 0.03 (-0.15-0.27) | 10.64 (9.17-13.70) | 100 |
| $p_0$ | -0.01 (-0.10-0.10) | 5.52 (5.15-6.09) | 100 | 0.00 (-0.10-0.11) | 5.57 (5.09-5.96) | 100 |
| $\sigma$ | 0.00 (-0.04-0.05) | 2.60 (2.38-2.90) | 100 | 0.00 (-0.05-0.04) | 2.58 (2.32-2.83) | 100 |

Table 6: Relative bias (RB) and coefficient of variation (CV) for the 100 simulated datasets for the scenario with $\beta_h = 1$, $\beta_w = 1$, and for a) $N_1 = 120$ and b) $N_1 = 250$. Values within brackets show the 95% quantiles.

(a) $N_1 = 120$

| | **Random** | | | **Gradient** | | |
|---|---|---|---|---|---|---|
| Parameters | RB | CV | Coverage | RB | CV | Coverage |
| | | | $\beta_h = 1$ $\beta_w = 1$ | | | |
| $N_1$ | 0.01 (-0.13-0.18) | 10.26 (8.91-12.14) | 97 | 0.03 (-0.14-0.28) | 9.93 (8.54-11.10) | 90 |
| $N_2$ | 0.01 (-0.18-0.16) | 8.66 (7.56-9.94) | 95 | 0.01 (-0.13-0.16) | 8.36 (7.17-9.66) | 97 |
| $N_3$ | 0.02 (-0.12-0.19) | 8.20 (7.38-9.37) | 94 | 0.01 (-0.13-0.15) | 7.83 (6.86-8.76) | 95 |
| $N_4$ | 0.01 (-0.15-0.14) | 8.29 (7.31-9.49) | 98 | 0.01 (-0.11-0.15) | 7.73 (6.94-8.79) | 99 |
| $N_5$ | 0.00 (-0.14-0.18) | 9.24 (8.17-10.02) | 98 | 0.02 (-0.11-0.18) | 8.44 (7.60-9.56) | 96 |
| $\beta_h$ | -0.04 (-1.31-1.47) | 61.72 (33.12-565.20) | 100 | 0.05 (-0.44-0.60) | 24.15 (19.24-38.92) | 100 |
| $\beta_w$ | 0.18 (-1.03-1.65) | 83.17 (50.00-4032.25) | 100 | 0.10 (-0.68-1.19) | 38.86 (25.78-114.36) | 100 |
| $\tau$ | 0.01 (-0.13-0.14) | 7.22 (6.42-8.39) | 100 | 0.01 (-0.11-0.13) | 6.98 (6.03-8.05) | 100 |
| $mh_h$ | 0.05 (-0.12-0.30) | 9.76 (7.61-18.42) | 100 | 0.02 (-0.13-0.16) | 8.23 (7.28-9.45) | 100 |
| $mh_w$ | 0.10 (-0.23-0.55) | 22.37 (14.28-38.08) | 100 | 0.07 (-0.20-0.39) | 16.16 (13.30-23.20) | 100 |
| $p_0$ | 0.00 (-0.13-0.14) | 7.98 (7.33-8.96) | 100 | 0.01 (-0.13-0.16) | 7.76 (7.10-8.59) | 100 |
| $\sigma$ | 0.00 (-0.07-0.09) | 3.75 (3.39-4.23) | 100 | 0.00 (-0.07-0.06) | 3.63 (3.24-4.15) | 100 |

(b) $N_1 = 250$

| | **Random** | | | **Gradient** | | |
|---|---|---|---|---|---|---|
| Parameters | RB | CV | Coverage | RB | CV | Coverage |
| | | | $\beta_h = 1$ $\beta_w = 1$ | | | |
| $N_1$ | 0.02 (-0.11-0.16) | 7.02 (6.48-7.83) | 93 | 0.02 (-0.10-0.20) | 6.90 (6.26-7.46) | 92 |
| $N_2$ | 0.00 (-0.10-0.15) | 5.96 (5.50-6.56) | 94 | 0.03 (-0.07-0.14) | 5.81 (5.37-6.28) | 96 |
| $N_3$ | 0.02 (-0.08-0.12) | 5.64 (5.22-6.27) | 96 | 0.02 (-0.10-0.12) | 5.37 (4.94-5.72) | 93 |
| $N_4$ | 0.02 (-0.08-0.15) | 5.67 (5.07-6.21) | 93 | 0.02 (-0.09-0.14) | 5.25 (4.85-5.75) | 91 |
| $N_5$ | 0.02 (-0.13-0.15) | 6.25 (5.70-6.92) | 92 | 0.01 (-0.15-0.13) | 5.75 (5.14-6.32) | 89 |
| $\beta_h$ | 0.17 (-0.59-1.13) | 35.16 (24.21-112.59) | 100 | 0.00 (-0.32-0.42) | 17.00 (14.44-21.29) | 100 |
| $\beta_w$ | 0.14 (-1.02-1.50) | 51.53 (32.64-740.10) | 100 | 0.06 (-0.55-1.00) | 25.00 (18.37-48.81) | 100 |
| $\tau$ | 0.00 (-0.09-0.10) | 5.00 (4.43-5.49) | 100 | -0.01 (-0.09-0.08) | 4.83 (4.44-5.41) | 100 |
| $mh_h$ | 0.04 (-0.08-0.32) | 6.88 (5.28-12.92) | 100 | 0.01 (-0.10-0.12) | 5.61 (5.26-6.12) | 100 |
| $mh_w$ | 0.05 (-0.16-0.51) | 14.89 (9.44-26.22) | 100 | 0.01 (-0.15-0.36) | 10.44 (9.43-14.20) | 100 |
| $p_0$ | 0.00 (-0.10-0.12) | 5.45 (5.06-6.00) | 100 | 0.01 (-0.10-0.09) | 5.40 (5.08-5.71) | 100 |
| $\sigma$ | 0.00 (-0.05-0.05) | 2.55 (2.36-2.84) | 100 | 0.00 (-0.04-0.05) | 2.50 (2.34-2.69) | 100 |

#### 1.2 Not accounting for spatial heterogeneity in mortality

Table 7: Percentage of models converged, identifiable and, both converged and identifiable for the 100 simulated datasets of 16 scenarios tested when misspecifying the OPSCR model by not accounting for spatial heterogeneity in mortality.

| $N_1$ | $\beta_h$ | $\beta_w$ | Spatial Covariate | % Converged | % Identifiable | % Converged and Identifiable |
|---|---|---|---|---|---|---|
| 120.00 | 1.00 | 1.00 | Gradient | 100 | 99 | 99 |
| 120.00 | -1.00 | 1.00 | Gradient | 100 | 100 | 100 |
| 120.00 | 1.00 | -1.00 | Gradient | 100 | 100 | 100 |
| 120.00 | -1.00 | -1.00 | Gradient | 100 | 100 | 100 |
| 250.00 | 1.00 | 1.00 | Gradient | 100 | 100 | 100 |
| 250.00 | -1.00 | 1.00 | Gradient | 100 | 100 | 100 |
| 250.00 | 1.00 | -1.00 | Gradient | 100 | 100 | 100 |
| 250.00 | -1.00 | -1.00 | Gradient | 100 | 100 | 100 |
| 120.00 | 1.00 | 1.00 | Random | 100 | 99 | 99 |
| 120.00 | -1.00 | 1.00 | Random | 100 | 99 | 99 |
| 120.00 | 1.00 | -1.00 | Random | 100 | 100 | 100 |
| 120.00 | -1.00 | -1.00 | Random | 100 | 98 | 98 |
| 250.00 | 1.00 | 1.00 | Random | 100 | 100 | 100 |
| 250.00 | -1.00 | 1.00 | Random | 100 | 100 | 100 |
| 250.00 | 1.00 | -1.00 | Random | 100 | 100 | 100 |
| 250.00 | -1.00 | -1.00 | Random | 100 | 100 | 100 |

Table 8: Relative bias (RB) and coefficient of variation (CV) for the 100 simulated datasets of the scenario with $\beta_h$ = 1, $\beta_w$ = -1, and a) $N_1$ = 120 and b) $N_1$ = 250 when not accounting for spatial heterogeneity in mortality. * denotes parameters for which we reported the average parameter value instead of RB, as RB could not be computed due to model misspecification. Values within brackets show the 95% quantiles.

(a) $N_1 = 120$

| | **Random** | | | **Gradient** | | |
|---|---|---|---|---|---|---|
| Parameters | RB | CV | Coverage | RB | CV | Coverage |
| | | | $\beta_h = 1$ $\beta_w$ = -1 | | | |
| $N_1$ | 0.02 (-0.18-0.20) | 10.31 (8.67-11.83) | 95 | 0.07 (-0.14-0.24) | 9.82 (8.31-11.39) | 91 |
| $N_2$ | 0.02 (-0.13-0.20) | 8.74 (7.55-10.13) | 95 | 0.10 (-0.08-0.23) | 8.53 (7.66-9.64) | 88 |
| $N_3$ | 0.02 (-0.10-0.19) | 8.30 (7.28-9.30) | 94 | 0.11 (-0.05-0.27) | 8.26 (7.31-9.13) | 80 |
| $N_4$ | 0.02 (-0.11-0.20) | 8.35 (7.55-9.40) | 93 | 0.10 (-0.04-0.21) | 8.29 (7.48-9.25) | 91 |
| $N_5$ | 0.03 (-0.16-0.23) | 9.27 (8.33-10.63) | 94 | 0.08 (-0.08-0.23) | 9.55 (8.67-10.90) | 94 |
| $\beta_h$ | - | - | - | - | - | - |
| $\beta_w$ | - | - | - | - | - | - |
| $\tau$ | -0.01 (-0.16-0.11) | 7.32 (6.38-9.37) | 94 | -0.02 (-0.14-0.14) | 7.36 (6.39-8.86) | 96 |
| $mh_h$* | -2.13 (-2.46–1.78) | 7.38 (6.82-8.50) | - | -1.90 (-2.16–1.65) | 7.68 (7.00-8.65) | - |
| $mh_w$* | -1.65 (-2.17–1.14) | 13.69 (12.25-16.53) | - | -1.56 (-2.00–1.24) | 13.43 (12.27-15.07) | - |
| $p_0$ | 0.01 (-0.13-0.15) | 7.97 (7.35-8.99) | 97 | 0.01 (-0.15-0.15) | 8.08 (7.27-8.86) | 98 |
| $\sigma$ | -0.01 (-0.07-0.07) | 3.72 (3.36-4.31) | 95 | 0.00 (-0.08-0.07) | 3.73 (3.35-4.17) | 95 |

(b) $N_1 = 250$

| | **Random** | | | **Gradient** | | |
|---|---|---|---|---|---|---|
| Parameters | RB | CV | Coverage | RB | CV | Coverage |
| | | | $\beta_h = 1$ $\beta_w$ = -1 | | | |
| $N_1$ | 0.01 (-0.11-0.17) | 7.09 (6.45-7.68) | 93 | 0.06 (-0.07-0.18) | 6.82 (6.33-7.44) | 88 |
| $N_2$ | 0.02 (-0.08-0.16) | 6.00 (5.50-6.46) | 94 | 0.07 (-0.02-0.18) | 5.96 (5.53-6.47) | 76 |
| $N_3$ | 0.00 (-0.09-0.12) | 5.62 (5.25-6.30) | 94 | 0.07 (-0.05-0.19) | 5.74 (5.35-6.28) | 74 |
| $N_4$ | 0.03 (-0.07-0.13) | 5.65 (5.28-6.17) | 96 | 0.07 (-0.03-0.19) | 5.79 (5.28-6.30) | 75 |
| $N_5$ | 0.02 (-0.11-0.13) | 6.36 (5.91-6.95) | 98 | 0.05 (-0.08-0.19) | 6.68 (6.11-7.26) | 87 |
| $\beta_h$ | - | - | - | - | - | - |
| $\beta_w$ | - | - | - | - | - | - |
| $\tau$ | 0.00 (-0.09-0.10) | 4.97 (4.39-5.67) | 94 | -0.02 (-0.12-0.08) | 5.12 (4.57-5.79) | 93 |
| $mh_h$* | -2.12 (-2.37–1.85) | 5.11 (4.82-5.58) | - | -1.88 (-2.12–1.74) | 5.34 (5.00-5.77) | - |
| $mh_w$* | -1.61 (-2.04–1.29) | 9.22 (8.55-9.93) | - | -1.51 (-1.83–1.30) | 9.22 (8.57-10.01) | - |
| $p_0$ | 0.01 (-0.07-0.12) | 5.50 (5.15-5.93) | 95 | -0.02 (-0.10-0.14) | 5.63 (5.27-6.13) | 93 |
| $\sigma$ | -0.01 (-0.05-0.05) | 2.55 (2.39-2.77) | 94 | 0.00 (-0.06-0.05) | 2.61 (2.39-2.85) | 89 |

Table 9: Relative bias (RB) and coefficient of variation (CV) for the 100 simulated datasets of the scenario with $\beta_h$ = -1, $\beta_w$ = -1, and a) $N_1 = 120$ and b) $N_1 = 250$ when not accounting for spatial heterogeneity in mortality. * denotes parameters for which we reported the average parameter value instead of RB, as RB could not be computed due to model misspecification. Values within brackets show the 95% quantiles.

(a) $N_1 = 120$

| | Random | | | Gradient | | |
|---|---|---|---|---|---|---|
| Parameters | RB | CV | Coverage | RB | CV | Coverage |
| | | | $\beta_h$ = -1 $\beta_w$ = -1 | | | |
| $N_1$ | 0.03 (-0.15-0.20) | 10.21 (8.78-11.59) | 98 | 0.04 (-0.15-0.22) | 9.94 (8.49-11.71) | 95 |
| $N_2$ | 0.04 (-0.18-0.20) | 8.54 (7.64-10.12) | 92 | 0.02 (-0.15-0.21) | 8.75 (7.67-9.84) | 94 |
| $N_3$ | 0.02 (-0.12-0.20) | 8.15 (7.23-9.18) | 92 | 0.02 (-0.16-0.14) | 8.33 (7.38-9.76) | 97 |
| $N_4$ | 0.02 (-0.13-0.18) | 8.20 (7.17-9.30) | 94 | -0.01 (-0.16-0.16) | 8.22 (7.50-9.35) | 95 |
| $N_5$ | 0.04 (-0.17-0.20) | 9.03 (8.05-10.23) | 94 | -0.01 (-0.20-0.16) | 9.03 (8.24-10.03) | 95 |
| $\beta_h$ | - | - | - | - | - | - |
| $\beta_w$ | - | - | - | - | - | - |
| $\tau$ | -0.01 (-0.18-0.17) | 7.11 (6.25-8.80) | 88 | 0.00 (-0.15-0.12) | 7.21 (6.20-9.05) | 95 |
| $mh_h$* | -2.22 (-2.55–1.89) | 7.25 (6.69-8.23) | - | -2.20 (-2.51–1.91) | 7.34 (6.76-8.32) | - |
| $mh_w$* | -1.70 (-2.23–1.27) | 13.65 (12.27-16.21) | - | -1.65 (-2.13–1.24) | 13.71 (12.02-15.69) | - |
| $p_0$ | 0.01 (-0.14-0.14) | 7.84 (7.16-8.78) | 96 | 0.00 (-0.14-0.15) | 7.89 (7.21-8.86) | 97 |
| $\sigma$ | -0.01 (-0.07-0.06) | 3.68 (3.33-4.14) | 96 | 0.01 (-0.06-0.07) | 3.75 (3.32-4.25) | 96 |

(b) $N_1 = 250$

| | Random | | | Gradient | | |
|---|---|---|---|---|---|---|
| Parameters | RB | CV | Coverage | RB | CV | Coverage |
| | | | $\beta_h$ = -1 $\beta_w$ = -1 | | | |
| $N_1$ | 0.02 (-0.13-0.15) | 7.05 (6.48-7.88) | 94 | 0.04 (-0.09-0.22) | 7.02 (6.21-7.64) | 88 |
| $N_2$ | 0.01 (-0.12-0.13) | 5.97 (5.47-6.56) | 93 | 0.03 (-0.08-0.14) | 6.09 (5.65-6.67) | 94 |
| $N_3$ | 0.01 (-0.11-0.11) | 5.64 (5.25-6.27) | 95 | 0.01 (-0.09-0.15) | 5.76 (5.35-6.21) | 93 |
| $N_4$ | 0.01 (-0.10-0.12) | 5.59 (5.09-6.17) | 94 | -0.02 (-0.14-0.11) | 5.72 (5.25-6.23) | 90 |
| $N_5$ | 0.00 (-0.11-0.15) | 6.27 (5.63-6.95) | 92 | -0.04 (-0.14-0.08) | 6.26 (5.77-6.83) | 94 |
| $\beta_h$ | - | - | - | - | - | - |
| $\beta_w$ | - | - | - | - | - | - |
| $\tau$ | -0.01 (-0.10-0.09) | 4.97 (4.42-5.84) | 94 | 0.01 (-0.09-0.11) | 5.04 (4.55-6.02) | 92 |
| $mh_h$* | -2.19 (-2.43–1.91) | 5.05 (4.73-5.54) | - | -2.20 (-2.42–1.98) | 5.10 (4.81-5.50) | - |
| $mh_w$* | -1.65 (-2.04–1.29) | 9.24 (8.56-9.99) | - | -1.62 (-1.97–1.32) | 9.28 (8.58-9.94) | - |
| $p_0$ | 0.00 (-0.09-0.13) | 5.47 (5.02-5.87) | 95 | -0.01 (-0.12-0.07) | 5.53 (5.20-6.05) | 91 |
| $\sigma$ | 0.00 (-0.04-0.05) | 2.57 (2.33-2.78) | 98 | 0.00 (-0.05-0.05) | 2.59 (2.41-2.85) | 93 |

Table 10: Relative bias (RB) and coefficient of variation (CV) for the 100 simulated datasets of the scenario with $\beta_h$ = -1, $\beta_w$ = 1, and a) $N_1$ = 120 and b) $N_1$ = 250 when not accounting for spatial heterogeneity in mortality. * denotes parameters for which we reported the average parameter value instead of RB, as RB could not be computed due to model misspecification. Values within brackets show the 95% quantiles.

(a) $N_1 = 120$

| | Random | | | Gradient | | |
|---|---|---|---|---|---|---|
| Parameters | RB | CV | Coverage | RB | CV | Coverage |
| | | | $\beta_h$ = -1 $\beta_w = 1$ | | | |
| $N_1$ | 0.04 (-0.14-0.18) | 10.10 (8.81-11.77) | 100 | 0.04 (-0.12-0.17) | 9.75 (8.59-10.93) | 99 |
| $N_2$ | 0.03 (-0.10-0.16) | 8.55 (7.48-9.60) | 98 | 0.06 (-0.08-0.22) | 8.55 (7.40-9.92) | 92 |
| $N_3$ | 0.04 (-0.15-0.16) | 8.15 (7.18-9.38) | 95 | 0.07 (-0.10-0.20) | 8.31 (7.37-9.24) | 96 |
| $N_4$ | 0.03 (-0.13-0.20) | 8.34 (7.17-9.51) | 95 | 0.07 (-0.10-0.21) | 8.47 (7.51-9.47) | 89 |
| $N_5$ | 0.01 (-0.19-0.20) | 9.28 (8.50-10.68) | 94 | 0.08 (-0.13-0.27) | 9.63 (8.50-10.74) | 91 |
| $\beta_h$ | - | - | - | - | - | - |
| $\beta_w$ | - | - | - | - | - | - |
| $\tau$ | 0.00 (-0.15-0.13) | 7.17 (6.39-8.61) | 95 | -0.01 (-0.16-0.13) | 7.32 (6.19-8.78) | 91 |
| $mh_h$* | -2.12 (-2.52–1.81) | 7.37 (6.83-8.32) | - | -1.91 (-2.11–1.63) | 7.77 (7.09-8.90) | - |
| $mh_w$* | -1.61 (-2.13–1.18) | 13.66 (12.25-15.56) | - | -1.65 (-2.22–1.25) | 13.81 (12.39-16.39) | - |
| $p_0$ | -0.02 (-0.12-0.13) | 7.98 (7.24-9.04) | 98 | 0.00 (-0.16-0.19) | 8.12 (7.27-8.97) | 92 |
| $\sigma$ | 0.01 (-0.06-0.07) | 3.76 (3.36-4.27) | 97 | 0.00 (-0.08-0.08) | 3.78 (3.30-4.25) | 94 |

(b) $N_1 = 250$

| | Random | | | Gradient | | |
|---|---|---|---|---|---|---|
| Parameters | RB | CV | Coverage | RB | CV | Coverage |
| | | | $\beta_h$ = -1 $\beta_w = 1$ | | | |
| $N_1$ | 0.03 (-0.10-0.14) | 7.04 (6.57-7.76) | 98 | 0.04 (-0.10-0.16) | 6.72 (6.12-7.32) | 91 |
| $N_2$ | 0.03 (-0.11-0.16) | 6.02 (5.48-6.78) | 93 | 0.05 (-0.05-0.17) | 5.84 (5.28-6.40) | 87 |
| $N_3$ | 0.03 (-0.09-0.13) | 5.64 (5.20-6.29) | 93 | 0.07 (-0.05-0.17) | 5.66 (5.09-6.03) | 83 |
| $N_4$ | 0.02 (-0.08-0.14) | 5.65 (5.26-6.31) | 91 | 0.07 (-0.05-0.19) | 5.74 (5.22-6.21) | 83 |
| $N_5$ | 0.02 (-0.11-0.15) | 6.39 (5.78-6.94) | 93 | 0.06 (-0.04-0.17) | 6.53 (6.13-7.07) | 84 |
| $\beta_h$ | - | - | - | - | - | - |
| $\beta_w$ | - | - | - | - | - | - |
| $\tau$ | 0.00 (-0.11-0.10) | 5.04 (4.39-5.83) | 91 | -0.01 (-0.10-0.09) | 5.03 (4.44-5.79) | 94 |
| $mh_h$* | -2.11 (-2.43–1.82) | 5.15 (4.85-5.56) | - | -1.91 (-2.08–1.69) | 5.33 (4.95-5.84) | - |
| $mh_w$* | -1.60 (-2.06–1.27) | 9.25 (8.53-10.16) | - | -1.67 (-1.97–1.42) | 9.28 (8.55-10.04) | - |
| $p_0$ | -0.02 (-0.10-0.10) | 5.52 (5.12-6.14) | 96 | 0.00 (-0.10-0.11) | 5.54 (5.07-5.93) | 94 |
| $\sigma$ | 0.00 (-0.05-0.05) | 2.59 (2.37-2.88) | 96 | 0.00 (-0.05-0.04) | 2.57 (2.35-2.81) | 98 |

Table 11: Relative bias (RB) and coefficient of variation (CV) for the 100 simulated datasets of the scenario with $\beta_h = 1$, $\beta_w = 1$, and a) $N_1 = 120$ and b) $N_1 = 250$ when not accounting for spatial heterogeneity in mortality. * denotes parameters for which we reported the average parameter value (95% quantiles) instead of RB, as RB could not be computed due to model misspecification. Values within brackets show the 95% quantiles.

(a) $N_1 = 120$

| | **Random** | | | **Gradient** | | |
|---|---|---|---|---|---|---|
| Parameters | RB | CV | Coverage | RB | CV | Coverage |
| | | | $\beta_h = 1$ $\beta_w = 1$ | | | |
| $N_1$ | 0.02 (-0.15-0.20) | 10.22 (8.83-12.21) | 98 | 0.07 (-0.10-0.32) | 9.90 (8.33-10.95) | 86 |
| $N_2$ | 0.02 (-0.14-0.17) | 8.61 (7.66-9.89) | 95 | 0.06 (-0.07-0.21) | 8.47 (7.21-9.73) | 93 |
| $N_3$ | 0.03 (-0.13-0.19) | 8.30 (7.28-9.31) | 95 | 0.04 (-0.09-0.19) | 8.06 (7.18-9.06) | 93 |
| $N_4$ | 0.01 (-0.15-0.20) | 8.33 (7.32-9.38) | 91 | 0.03 (-0.11-0.19) | 8.07 (7.32-9.07) | 95 |
| $N_5$ | 0.01 (-0.15-0.19) | 9.20 (8.13-10.07) | 96 | 0.02 (-0.11-0.18) | 8.92 (8.21-9.86) | 98 |
| $\beta_h$ | - | - | - | - | - | - |
| $\beta_w$ | - | - | - | - | - | - |
| $\tau$ | 0.00 (-0.13-0.13) | 7.09 (6.15-8.71) | 97 | 0.00 (-0.12-0.13) | 7.00 (6.11-8.16) | 98 |
| $mh_h$* | -2.17 (-2.49–1.84) | 7.39 (6.72-8.25) | - | -2.23 (-2.54–1.90) | 7.20 (6.67-7.94) | - |
| $mh_w$* | -1.66 (-2.26–1.27) | 13.73 (12.43-16.53) | - | -1.81 (-2.19–1.44) | 13.50 (12.19-16.26) | - |
| $p_0$ | 0.00 (-0.14-0.14) | 8.01 (7.16-8.84) | 98 | 0.01 (-0.13-0.17) | 7.76 (7.15-8.62) | 95 |
| $\sigma$ | -0.01 (-0.07-0.08) | 3.76 (3.31-4.18) | 94 | 0.00 (-0.07-0.06) | 3.62 (3.25-4.15) | 94 |

(b) $N_1 = 250$

| | **Random** | | | **Gradient** | | |
|---|---|---|---|---|---|---|
| Parameters | RB | CV | Coverage | RB | CV | Coverage |
| | | | $\beta_h = 1$ $\beta_w = 1$ | | | |
| $N_1$ | 0.03 (-0.12-0.16) | 7.02 (6.35-7.85) | 95 | 0.06 (-0.06-0.23) | 6.90 (6.33-7.43) | 87 |
| $N_2$ | 0.00 (-0.09-0.15) | 5.97 (5.46-6.59) | 94 | 0.07 (-0.02-0.18) | 5.90 (5.47-6.36) | 75 |
| $N_3$ | 0.01 (-0.09-0.13) | 5.65 (5.25-6.31) | 94 | 0.05 (-0.09-0.16) | 5.58 (5.17-5.97) | 82 |
| $N_4$ | 0.02 (-0.08-0.14) | 5.67 (5.09-6.21) | 95 | 0.04 (-0.07-0.19) | 5.57 (5.18-6.00) | 89 |
| $N_5$ | 0.01 (-0.14-0.15) | 6.25 (5.79-6.89) | 93 | 0.01 (-0.14-0.16) | 6.10 (5.65-6.63) | 88 |
| $\beta_h$ | - | - | - | - | - | - |
| $\beta_w$ | - | - | - | - | - | - |
| $\tau$ | 0.00 (-0.09-0.10) | 4.97 (4.43-5.67) | 95 | -0.01 (-0.09-0.08) | 4.81 (4.43-5.33) | 97 |
| $mh_h$* | -2.17 (-2.45–1.93) | 5.08 (4.57-5.50) | - | -2.23 (-2.42–2.03) | 4.96 (4.68-5.30) | - |
| $mh_w$* | -1.62 (-2.09–1.36) | 9.25 (8.48-10.18) | - | -1.79 (-2.22–1.53) | 9.08 (8.57-9.76) | - |
| $p_0$ | 0.01 (-0.10-0.12) | 5.46 (5.01-5.95) | 94 | 0.00 (-0.10-0.10) | 5.40 (5.08-5.70) | 97 |
| $\sigma$ | 0.00 (-0.05-0.05) | 2.56 (2.34-2.79) | 92 | 0.00 (-0.04-0.05) | 2.52 (2.34-2.68) | 95 |

### 2 Appendix 2. Density dependence in survival

Table 12: Summary of the simulation with an effect of local density on survival. The average number (and standard deviation), across all 100 datasets, of individuals considered alive ($z_i = 2$), and detected alive for all consecutive occasions.

| $N_1$ | $\beta_{phi}$ | $\beta_{phi_0}$ | **Alive** | | | | | **Detected** | | | | |
|---|---|---|---|---|---|---|---|---|---|---|---|---|
| $N_1$ | $\beta_{phi}$ | $\beta_{phi_0}$ | 1 | 2 | 3 | 4 | 5 | 1 | 2 | 3 | 4 | 5 |
| 250 | -1 | 1.6 | 253.2(11.6) | 223.8(10.4) | 213.0(10.4) | 207.9(11.2) | 206.0(10.6) | 149.4(10.8) | 136.5(10.3) | 131.2( 8.8) | 129.9( 9.9) | 131.2( 9.5) |

Table 13: Percentage of models converged, identifiable and both converged and identifiable for the 100 simulated datasets. a)"With" and b)"Without" represent results obtained when fitting an OPSCR model that accounted and that did not account for the effect of density on survival, respectively. Note that both models were fitted to the same 100 replicated datasets.

(a) *With*

| $N_1$ | $\beta_{phi}$ | $\beta_{phi_0}$ | % Converged | % Identifiable | % Converged and Identifiable |
|---|---|---|---|---|---|
| 250.00 | -1.00 | 1.60 | 100 | 86 | 86 |

(b) *Without*

| [redacted] | $\beta_{phi}$ | $\beta_{phi_0}$ | % Converged | % Identifiable | % Converged and Identifiable |
|---|---|---|---|---|---|
| 250.00 | -1.00 | 1.60 | 100 | 100 | 100 |

Table 14: Relative bias (RB), standard deviation (SD) and coverage for the 100 replicated datasets of the local effect of density on survival. "With" and "Without" represent results obtained when fitting an OPSCR model that accounted and that did not account for the effect of density on survival, respectively. Note that the same 100 data sets was fitted to both models. * Indicates estimates for which RB could not be computed because the model was misspecified. Values within brackets show the 95% quantiles.

| [redacted] | **With** | | | **Without** | | |
|---|---|---|---|---|---|---|
| Parameters | RB | SD | Coverage | RB | SD | Coverage |
| $N_1$ | 0.01 (-0.10-0.11) | 13.49 (12.43-14.64) | 92 | 0.01 (-0.10-0.12) | 13.19 (12.16-14.61) | 92 |
| $N_2$ | 0.00 (-0.08-0.08) | 10.45 (9.61-11.44) | 99 | 0.01 (-0.06-0.10) | 10.60 (9.84-11.51) | 97 |
| $N_3$ | 0.00 (-0.08-0.09) | 9.81 (8.92-10.64) | 97 | 0.00 (-0.06-0.09) | 9.90 (9.07-10.68) | 96 |
| $N_4$ | 0.00 (-0.06-0.09) | 9.73 (8.87-10.81) | 97 | 0.00 (-0.07-0.09) | 9.78 (8.99-10.71) | 98 |
| $N_5$ | 0.01 (-0.06-0.12) | 10.09 (9.12-11.06) | 92 | 0.00 (-0.07-0.11) | 10.19 (9.33-11.35) | 96 |
| $\beta_\phi$ | 0.13 (-0.63-0.82) | 0.43 (0.32-0.61) | 96 | - | - | - |
| $\tau$ | 0.00 (-0.08-0.06) | 0.03 (0.02-0.03) | 98 | 0.00 (-0.08-0.06) | 0.03 (0.02-0.03) | 98 |
| $p_0$ | 0.00 (-0.10-0.07) | 0.00 (0.00-0.00) | 94 | 0.00 (-0.10-0.07) | 0.00 (0.00-0.01) | 94 |
| $\phi_0$ | 0.13 (-0.70-0.83) | 0.73 (0.53-1.04) | 96 | -0.05 (-0.17-0.05)* | 0.06 (0.06-0.06) | - |
| $\sigma$ | 0.00 (-0.03-0.05) | 0.01 (0.01-0.01) | 98 | 0.00 (-0.03-0.05) | 0.01 (0.01-0.01) | 96 |

### 3 Appendix 3. Mapping spatial heterogeneity in mortality

We quantified the overall deviation between simulated and OPSCR model predicted spatial maps of mortality. With Bayesian models, we can plot 1) the predicted spatial pattern of $w$ and $h$ (using eqn (3) and (4) in the main text), but also 2) the realized $w$ and $h$ for each habitat cell using the individual posterior $z$ and $s$ locations. We computed the average predictions of $w$ and $h$ for both methods for each cell $r$ of the spatial domain $S$ and then computed the cell by cell relative error between the simulated and predicted maps. Cell relative error was averaged to obtain a single measure of overall relative error for each replicated dataset. Because we did not consider time-dependence in mortality, we only obtained one relative error measure when using the predicted spatial pattern from $w$ and $h$ (1)). However, we computed the relative error for each transition (4 in total) for the realized $w$ and $h$ (2)) since we had access to all four survival transitions using the individual posterior $z$ and $s$. We performed the calculation described above only for two scenarios with small ($N_1 = 120$) and large ($N_1 = 250$) population size, with a gradient deterministic spatial covariate, $\beta_w = 1$ and $\beta_h = -1$. For computational reasons, we thinned $z$ and $s$ by 10, which resulted in a total of 8400 posterior samples (i.e. 2800 samples per chain).

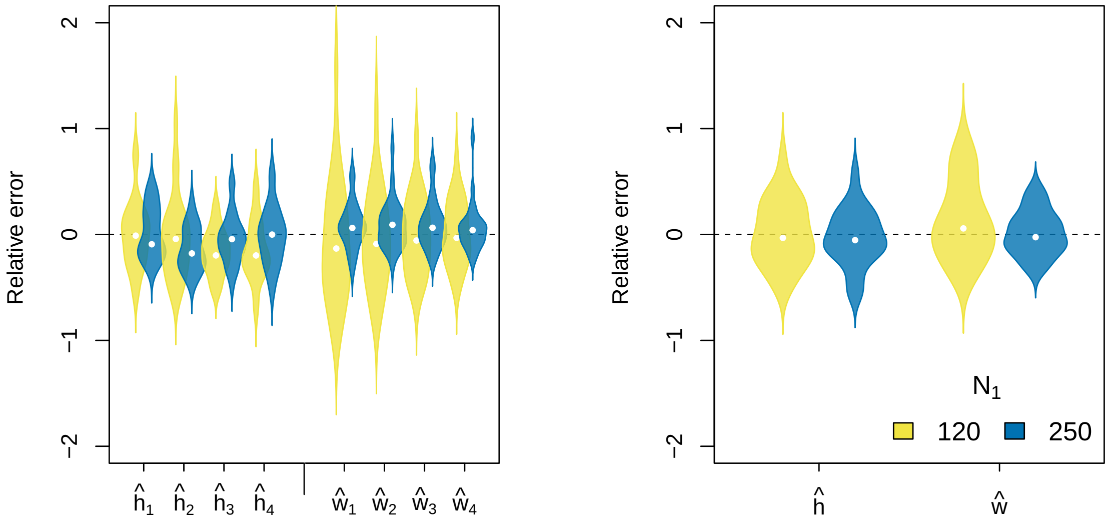


Figure 1: Violin plots of the distribution of the relative error in the spatial pattern of cause-specific mortality probabilities ($\hat{h}$ and $\hat{w}$) obtained for each transition from the posterior $s$ and $z$ (left) and the predicted $\hat{h}$ and $\hat{w}$ (right). Results are presented for only two scenarios with small ($N_1 = 120$) and large ($N_1 = 250$) population size, with a gradient spatial covariate, $\beta_w = 1$ and $\beta_h = -1$.

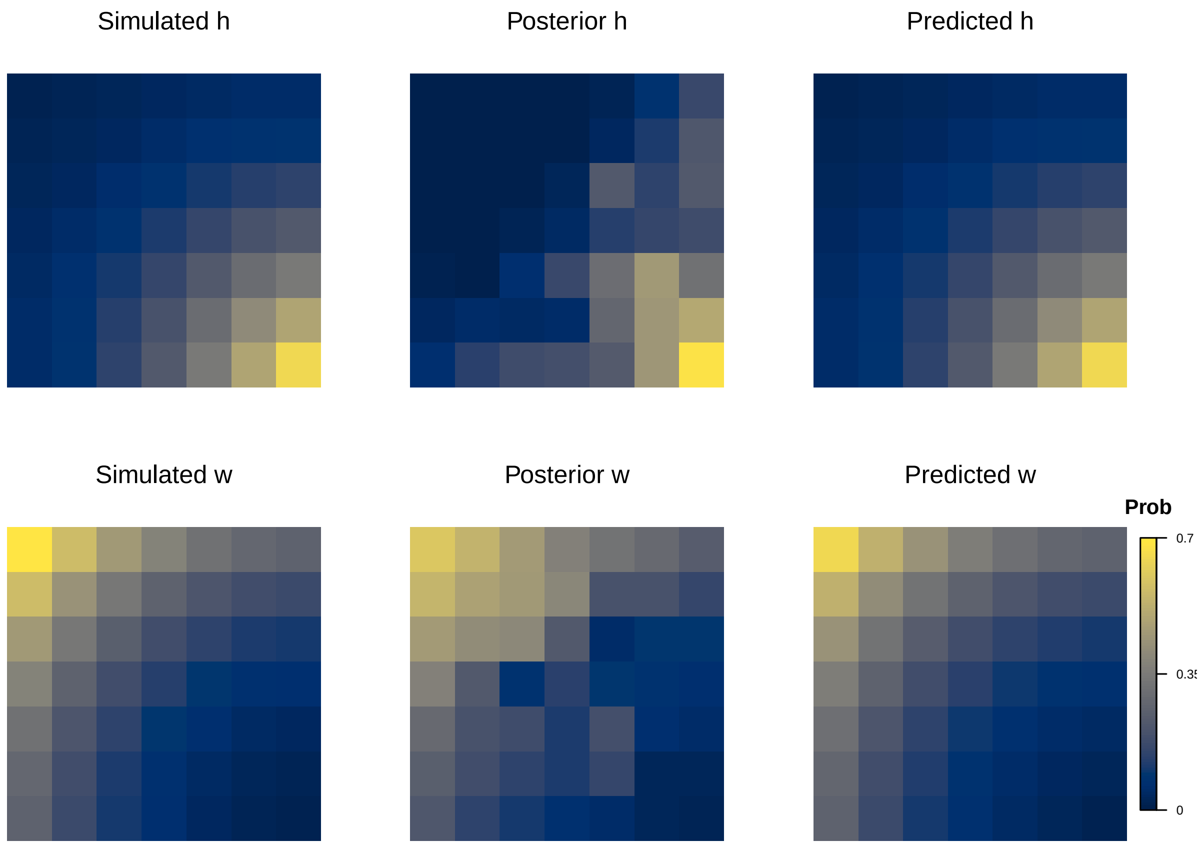


Figure 2: Example of the spatial pattern in mortality for $h$ (top row) and $w$ (bottom row) used to simulate the data (Simulated), estimated by the model using the average individual posterior $s$ and $z$ (Posterior), and predicted by the model using eqn 3 and 4 (Predicted). Note that $w$ and $h$ are on the probability (Prob) scale. Maps were computed using one replicated dataset and the second survival transition for the 'Posterior' maps.
