## Appendix 4 for "Estimating spatially variable and density-dependent survival using open-population spatial capture-recapture models"

Appendix 4: Using nimbleSCR to simulate from and fit Bayesian OPSCR models with spatial survival


### Appendix 4: Using nimbleSCR to simulate from and fit Bayesian OPSCR models with spatial survival

###### Cyril Milleret

#### 2022-02-25

In this vignette, we demonstrate how to use the nimbleSCR (Bischof et al. 2020) and NIMBLE packages (de Valpine et al. 2017; NIMBLE Development Team 2020) to simulate open-population spatial capture-recapture (OPSCR) data and fit flexible and efficient Bayesian OPSCR models with spatially hetereogenous mortality. We assume that we have access to detections of individuals alive but also dead recovery locations (Dupont et al. 2021). The OPSCR model is parameterized using a hazard rate formulation and two competing cause of mortality. For one cause of mortality (e.g. culling), we assume that all mortality events are recovered with an associated death location. See details in Milleret et al….

```
## Load packages
library(nimble)
library(nimbleSCR)
library(basicMCMCplots)

source("C:/Personal_Cloud/OneDrive/Work/PublicCodesGit/Public/SpatialSurvivalOPSCR/sampler_categorical_general1.R")
source("C:/Personal_Cloud/OneDrive/Work/PublicCodesGit/Public/SpatialSurvivalOPSCR/dcatState2Dead.R")
```

```
## Registering the following user-provided distributions: dcatState2Dead
```

#### 1. Simulate SCR data

##### 1.1 Habitat and trapping grid

As an example, we create a \(80 \times 100\) habitat grid with a resolution of 10 for each dimension. On the habitat, we center a \(60 \times 80\) trapping grid with also a resolution of 10 for each dimension, leaving an untrapped perimeter (buffer) with a width of 20 distance units on each side of the grid.

```
## Create habitat grid
coordsHabitatGridCenter <- cbind(rep(seq(23, 8, by = -5), 4),
                                 sort(rep(seq(8, 23, by = 5), 4)))
colnames(coordsHabitatGridCenter) <- c("x","y")

## Create trap grid
coordsObsCenter <- cbind(rep(seq(8, 23, by = 1), 16),
                         sort(rep(seq(23, 8, by = -1), 16)))
colnames(coordsObsCenter) <- c("x","y")

## Plot check
plot(coordsHabitatGridCenter[,"y"] ~ coordsHabitatGridCenter[,"x"],
     xlim = c(5,25), ylim = c(5,25),
     pch = 1, cex = 1.5) 
points(coordsObsCenter[,"y"] ~ coordsObsCenter[,"x"], col="red", pch=16 ) 
par(xpd=TRUE)
legend(x = 7, y = 7,
       legend=c("Habitat window centers", "Observation window centers"),
       pt.cex = c(1.5,1),
       horiz = T,
       pch=c(1,16),
       col=c("black", "red"),
       bty = 'n')
```

```
habitatMask <- matrix(1, nrow = 4, ncol= 4, byrow = TRUE)
```

##### 1.2 Rescale coordinates

To implement the local evaluation approach when fitting the SCR model (see Milleret et al. (2019) and Turek et al. (2021) for further details), we need to rescale the habitat and trapping grid coordinates so that each habitat cell is of dimension \(1 \times 1\). We also need to identify the lower and upper coordinates of each habitat cell using the ‘getLowAndUpCoords’ function.

```
## Rescale coordinates
scaledObjects <- scaleCoordsToHabitatGrid(
  coordsData = coordsObsCenter,
  coordsHabitatGridCenter = coordsHabitatGridCenter)

## Get lower and upper cell coordinates
lowerAndUpperCoords <- getWindowCoords(
  scaledHabGridCenter = scaledObjects$coordsHabitatGridCenterScaled,
  scaledObsGridCenter = scaledObjects$coordsDataScaled,
  plot.check = F)
```

We also set up the objects necessary to perform the local evaluation for the live detection using the ‘getLocalObjects’ function. Special care should be taken when chosing ‘dmax’ relative to \(\sigma\) (Milleret et al. 2019). Here we are using a value \(>3\*\sigma\) (see below for the \(\sigma\) chosen.

```
trapLocal <- getLocalObjects(habitatMask = habitatMask,
                             coords = scaledObjects$coordsDataScaled,
                             dmax = 1,
                             resizeFactor = 1,
                             plot.check = TRUE
)
```

We create the local objects to perform the local evaluation for the dead recovery locations using the ‘getLocalObjects’ function. We assume that dead recoveries from one cause mortality are also detected within the buffer area. We therefore use the coordinates of the entire spatial domain (and not only the detector coordinates) to define the local objects.

```
deadObsLocal <- getLocalObjects(habitatMask = habitatMask,
                             coords = scaledObjects$coordsHabitatGridCenterScaled,
                             dmax = 2,
                             resizeFactor = 1,
                             plot.check = TRUE
)
```

##### 1.3 Define model code

Note that we do not provide ‘detNums’ and ‘detIndices’ arguments in the ‘dbinomLocal\_normal’ function as we wish to use this model to simulate data (see ‘?dbinomLocal\_normal’ for further details).

```
modelCode <- nimbleCode({
  ##--------------------------------------------------------------------------------------------
  ##-----------------------------## 
  ##------ SPATIAL PROCESS ------##  
  ##-----------------------------##
  tau ~ dgamma(0.001, 0.001)
  logHabIntensity[1:numHabWindows] <- mu[1:numHabWindows]
  sumHabInt <- log(sum(mu[1:numHabWindows]))
  ## FIRST YEAR 
  for(i in 1:M){
    s[i, 1:2,1] ~ dbernppAC(
      lowerCoords = lowerHabCoords[1:numHabWindows, 1:2],
      upperCoords = upperHabCoords[1:numHabWindows, 1:2],
      logIntensities = logHabIntensity[1:numHabWindows],
      logSumIntensity = sumHabInt,
      habitatGrid = habitatGrid[1:y.max,1:x.max],
      numGridRows =  y.max,
      numGridCols = x.max
    )
  }#i
  
  ## T>1 
  for(t in 2:n.years){
    for(i in 1:M){
      s[i, 1:2, t] ~ dbernppACmovement_normal(lowerCoords = lowerHabCoords[1:numHabWindows, 1:2]
                                                , upperCoords = upperHabCoords[1:numHabWindows, 1:2]
                                                , s = s[i, 1:2, t - 1]
                                                , sd = tau
                                                , baseIntensities = mu[1:numHabWindows]
                                                , habitatGrid =  habitatGrid[1:y.max,1:x.max]
                                                , numGridRows = y.max
                                                , numGridCols = x.max
                                                , numWindows= numHabWindows
      )
    }#i
  }#t
  
  ##--------------------------------------------------------------------------------------------
  ##-------------------------------## 
  ##----- DEMOGRAPHIC PROCESS -----## 
  ##-------------------------------##    
  ## INTERCEPT MORTALITY HR
  mhH ~ dunif(-10, 10)
  mhW ~ dunif(-10, 10)
  
  ## CAUSE SPECIFIC SLOPE MORTALITY 
  betaH ~ dunif(-10,10)
  betaW ~ dunif(-10,10)
  
  ## FIRST YEAR INCLUSION PARAMETER 
  gamma1 ~ dunif(0,1)
  omeg1[1:2] <- c(1-gamma1, gamma1)
  
  ## TIME SPECIFIC INCLUSION PARAMETER 
  for(t in 1:n.years1){
    gamma[t] ~ dunif(0,1)
  }#t
  
  ## SPATIAL VITAL RATES 
  ## RECRUITMENT CONSTANT OVER SPACE 
  # for(t in 1:n.years1){
  #   for(r in 1:numHabWindows){
  #     gamma2[r,t] <- gamma[t] 
  #   }
  # }
  
  ## SURVIVAL (CAUSE-SPECIFIC MORTALITY SPATIALLY EXPLICIT)
  mhH1[1:numHabWindows] <- exp(mhH + betaH*habCov[1:numHabWindows])
  mhW1[1:numHabWindows] <- exp(mhW + betaW*habCov[1:numHabWindows])
  
  # derive mortality and surival probabilities 
  phi[1:numHabWindows] <- exp(-(mhH1[1:numHabWindows] + mhW1[1:numHabWindows]))
  h[1:numHabWindows] <-  (1 - phi[1:numHabWindows])* (mhH1[1:numHabWindows]/(mhH1[1:numHabWindows] +
                         mhW1[1:numHabWindows]))
  w[1:numHabWindows] <-  (1 - phi[1:numHabWindows])* (mhW1[1:numHabWindows]/(mhH1[1:numHabWindows] +
                         mhW1[1:numHabWindows]))
  
  ## STATE TRANSITION 
  for(i in 1:M){ 
    z[i,1] ~ dcat(omeg1[1:2]) 
    for(t in 1:n.years1){
      z[i,t+1] ~ dcatState2Dead(   z = z[i,t]
                                 , gamma = gamma[t]
                                 , hSpatial =  h[1:numHabWindows]
                                 , wSpatial = w[1:numHabWindows]
                                 , phiSpatial = phi[1:numHabWindows]
                                 , s = s[i,1:2,t]
                                 , habitatGrid = habitatGrid[1:y.max,1:x.max])
    }#i                                 
  }#t 
  
  ##---------------------------------------------------------------------------------------------   
  ##-----------------------------##
  ##----- DETECTION PROCESS -----## 
  ##-----------------------------##
  sigma ~ dunif(0,15)
  p0 ~ dunif(0,1)
  
  for(t in 1:n.years){
    for (i in 1:M){
      
      ## ALIVE DETECTIONS 
       y[i, 1:lengthYCombined,t] ~ dbinomLocal_normal(size = trials[1:n.traps],
                                                 p0 = p0,
                                                 s = s[i,1:2,t],
                                                 sigma = sigma,
                                                 trapCoords = trapCoords[1:n.traps,1:2],
                                                 localTrapsIndices = trapIndex[1:n.cells,1:maxNBDets],
                                                 localTrapsNum = nTraps[1:n.cells],
                                                 resizeFactor = resizeFactor,
                                                 habitatGrid = habitatIDDet[1:y.maxDet,1:x.maxDet],
                                                 indicator = isAlive[i,t],
                                                 lengthYCombined = lengthYCombined)
      
      
      ## DEAD RECOVERY 
      y.dead[i, 1:3,t] ~ dbernppLocalDetection_normal(
          lowerCoords= lowerHabCoords[1:numHabWindows,1:2]
        , upperCoords= upperHabCoords[1:numHabWindows,1:2]
        , s = s[i,1:2,t]
        , sd = sigma
        , baseIntensities = detReco[1:numHabWindows]
        , habitatGridLocal=habitatGridLocal[1:y.max,1:x.max]
        , resizeFactor=resizeFactor
        , localObsWindowIndices = localObsWindowIndices[1:numHabWindows,1:maxNBDetsReco]
        , numLocalObsWindows = numLocalObsWindows[1:numHabWindows]
        , numWindows = numHabWindows
        , indicator = z[i,t]==3
      )
    }#i
  }#t
  
  ##---------------------------------------------------------------------------------------------                                       
  ##----------------------------------------## 
  ##---------- DERIVED PARAMETERS ----------##
  ##----------------------------------------##
  for(i in 1:M){ 
    isAlive[i,1] <- (z[i,1] == 2) 
    for(t in 1:n.years1){
      isAlive[i,t+1] <- (z[i,t+1] == 2) 
    }
  }
  for(t in 1:n.years){
    N[t] <- sum(isAlive[1:M,t])
  }#t
})
```

##### 1.4 Define parameter values to simulate

The model formulation uses data augmentation to derive N estimates (Royle and Dorazio 2012). We therefore need to choose the total number of individuals *M* (detected + augmented).

```
M <- 500

## ASSIGN SIMULATED VALUES 
p0 <- 0.2
sigma <- 0.4

#expected number of individuals alive (z=2)at t=1
n.individualsT1 <- 200
n.years <- 5
mhW <- -2
mhH <- -2

betaH <- 1
betaW <- -1
tau <- 1.2

recruitment <- 0.3  
## calculate the gamma 
gamma <- n.individualsT1/M
Recruit <- n.individualsT1*recruitment ## 40 % of recruitment 
NeverAlive <- n.individualsT1
  
for(t in 2:n.years){
    Navai  <- M- NeverAlive[t-1]
    gamma[t] <- Recruit/Navai 
    NeverAlive[t] <- NeverAlive[t-1] + Recruit
}
```

We create a spatial covariate with a horizontal gradient that will be used to explain heterogeneity in mortality.

```
habCov <- as.numeric(scale(lowerAndUpperCoords$lowerHabCoords[,2])[,1])
habCovImage <- lowerAndUpperCoords$habitatGrid
habCovImage[] <- habCov[lowerAndUpperCoords$habitatGrid]
image(habCovImage)
```

```
### plot expected surv/mortality prob as a function of covariate 
mhH1 <- exp(mhH + betaH*habCov)
mhW1 <- exp(mhW + betaW*habCov)
  
phi <- exp(-(mhH1 + mhW1))
h <-  (1-phi) * (mhH1/(mhH1 + mhW1))
w <-  (1-phi) * (mhW1/(mhH1 + mhW1))
  
plot(phi ~ habCov, ylim=c(0,1), type="b", ylab="Probability")
points(h ~ habCov, col="red", type="b")
points(w ~ habCov, col="blue", type="b")
legend("topright", legend=c("phi","h","w"), 
       col=c("black","red","blue"), lty=c(1,1,1))
```

When simulating detections using this formulation of the SCR model in NIMBLE, all the information about detections (where and how many) is stored in ‘y’ in that order (See ?dbinomlocal\_normal for more details.):

- ‘detNums’ (total number of individual detections),
- ‘x’ (number of individual detections at each trap),
- ‘detIndices’ (id of the trap at which detections occur).

We now need to provide the maximum number of spatial recaptures that can be simulated per individual. We recommend using ‘trapLocal$numlocalindicesmax’ that defines the maximum number of traps available for detections when local evaluation is used. This will enable the simulation of as many spatial detections as allowed by the restrictions imposed by the local evaluation (defined by the ‘dmax’ argument from ‘getLocalObjects’). This means that the length of the ‘y’ observation vector for each individual is equal to the length of \(c(detNums, x, detIndices)\) and is therefore equal to \(lengthYCombined = 1+ trapLocal\$numLocalIndicesMax \* 2\).

```
lengthYCombined <- 1 + trapLocal$numLocalIndicesMax*2
```

##### 1.5 Create data, constants and inits objects

```
nimConstants <- list(M = M,
                     n.years = n.years,
                     n.years1 = n.years-1,
                     n.traps = dim(scaledObjects$coordsDataScaled)[1],
                     y.max = dim(habitatMask)[1],
                     x.max = dim(habitatMask)[2],
                     y.maxDet = dim(trapLocal$habitatGrid)[1],
                     x.maxDet = dim(trapLocal$habitatGrid)[2],
                     ResizeFactor = trapLocal$resizeFactor,
                     n.cells = dim(trapLocal$localIndices)[1],
                     maxNBDets = trapLocal$numLocalIndicesMax,
                     trapIndex = trapLocal$localIndices,
                     nTraps = trapLocal$numLocalIndices,
                     habitatIDDet = trapLocal$habitatGrid,
                     lengthYCombined = lengthYCombined,
                     numHabWindows = dim(lowerAndUpperCoords$lowerHabCoords)[1],
                     maxNBDetsReco = deadObsLocal$numLocalIndicesMax)


nimData <- list(trapCoords = scaledObjects$coordsDataScaled,
                trials = rep(1, dim(scaledObjects$coordsDataScaled)[1]),
                lowerHabCoords = lowerAndUpperCoords$lowerHabCoords,
                upperHabCoords = lowerAndUpperCoords$upperHabCoords,
                habitatGrid = lowerAndUpperCoords$habitatGrid,
                detReco = rep(1, dim(lowerAndUpperCoords$lowerHabCoords)[1]),
                habCov = habCov,
                mu = rep(1, dim(lowerAndUpperCoords$lowerHabCoords)[1]),
                habitatGridLocal = deadObsLocal$habitatGrid,
                localObsWindowIndices = deadObsLocal$localIndices,
                numLocalObsWindows = deadObsLocal$numLocalIndices,
                resizeFactor = deadObsLocal$resizeFactor
                )

# We set the parameter values as inits
nimInits <- list(sigma = sigma,
                 p0=p0,
                 tau=tau,
                 gamma = gamma[2:n.years],
                 gamma1 = gamma[1],
                 mhW = mhW,
                 mhH = mhH,
                 betaH = betaH,
                 betaW = betaW)
```

##### 1.6 Create NIMBLE model

```
model <- nimbleModel( code = modelCode,
                      constants = nimConstants,
                      data = nimData,
                      inits = nimInits,
                      check = F,       
                      calculate = F)
```

##### 1.7 Simualte SCR data from the NIMBLE model

We first need to obtain the list of nodes that will be simulated. We used the ‘getDependencies’ function from NIMBLE. Using the ‘simulate’ function from NIMBLE, we will then simulate the activity center (AC) locations (‘s’), the state of the individual (‘z’) and SCR observation data (‘y’) given the values we provided for ‘p0,’ ‘sigma’ and ‘psi.’

```
# FIRST WE GET THE NODES TO SIMULATE
#nodesToSim <- model$getDependencies(c("s", "z"), self=T)
nodesToSim <- model$getDependencies(c("p0","sigma","mhW", "mhH","mu",
                                      "gamma","tau","gamma1","betaH",
                                       "betaW"),
                                      self = F,
                                      downstream = T,
                                      returnScalarComponents = TRUE)
# THEN WE SIMULATE THOSE NODES 
set.seed(1)
model$simulate(nodesToSim, includeData = FALSE)

  # 
# 
# t=1
# i=1
#                          rcatState2Dead(n=1
#                                  , 
#                                  zit = model$z[i,t]
#                                  , 
#                                  gamma = model$gamma[t]#gamma2[1:numHabWindows,t]
#                                  , 
#                                  gammaVec = model$s[i,1:2,t]
#                                  , 
#                                  h = -999
#                                  , 
#                                  hVec =  model$h[1:numHabWindows]
#                                  , 
#                                  w = -999
#                                  , 
#                                  wVec = model$w[1:numHabWindows]
#                                  , 
#                                  phi = -999
#                                  , 
#                                  phiVec = model$phi[1:numHabWindows]
#                                  , 
#                                  s = model$s[i,1:2,t]
#                                  , 
#                                  habIndex = model$habitatGrid[1:y.max,1:x.max]
#                                  )
#
```

After running ‘simulate,’ the simulated data are stored in the ‘model’ object. For example, we can access the simulated ‘z’ and check how many individuals were considered alive and recovered at each time step :

```
N <- apply(model$z,2,function(x)sum(x==2))
N
```

```
## [1] 188 185 183 184 193
```

```
N.recoveredDead <- apply(model$z,2,function(x)sum(x==3))
N.recoveredDead
```

```
## [1]  0 24 36 38 31
```

#### 2. RUN MCMC WITH NIMBLE

Here, we build the NIMBLE model again using the simulated ‘y’ and ‘y.dead’ as data. For simplicity, we used the simulated ‘z’ as initial values. Then we can fit the SCR model with the simulated ‘y’ and ‘y.dead’ data set.

```
myZ <- model$z
z <- zInits <- model$z
whichDet <- apply(model$y, 3, function(x) x[,1]>0 )
whichNotDet <- apply(model$y, 3, function(x) x[,1]==0 )

# give NAS to individuals not detected
z[whichNotDet] <- NA
zInits[whichDet] <- NA

nimData$y <- model$y
nimData$y.dead <- model$y.dead

nimData$z <- z
nimInits$z <- zInits
nimInits$s <- model$s

# CREATE AND COMPILE THE NIMBLE MODEL
model <- nimbleModel( code = modelCode,
                      constants = nimConstants,
                      data = nimData,
                      inits = nimInits,
                      check = F,
                      calculate = F)
model$calculate()
```

```
## [1] -15865.97
```

```
cmodel <- compileNimble(model)
cmodel$calculate()
MCMCconf <- configureMCMC(model = model,
                          monitors = c("p0","sigma","mhW","mhH", "N",
                                         "gamma","tau","betaH","betaW"),
                          control = list(reflective = TRUE),
                          thin = 1)
## WE NEED TO SET THE NODES FOR WHICH dcatState2Dead IS USED AS A CATEGORICAL SAMPLER
###FIND WHICH Z NODES NEED TO BE REPLACED BY THE NEW SAMPLER. 
samplerConfList <- unlist(lapply(MCMCconf$getSamplers(),function(x) x$target))
zNodes <- samplerConfList[grep("z",samplerConfList)]
#find z nodes from the first year
zNodes <- zNodes[-grep( ", 1]",zNodes)]

MCMCconf
#remove samplers
MCMCconf$removeSamplers(zNodes)
for(i in 1:length(zNodes)){
  MCMCconf$addSampler(target = zNodes[i], type = 'sampler_categorical_general1',
                                  control=list("numCategories"= 4))
}

MCMC <- buildMCMC(MCMCconf)
cMCMC <- compileNimble(MCMC, project = model, resetFunctions = TRUE)
# RUN THE MCMC 
MCMCRuntime <- system.time(myNimbleOutput <- runMCMC( mcmc = cMCMC,
                                                             nburnin = 1000,
                                                             niter = 5000,
                                                             nchains = 3,
                                                             samplesAsCodaMCMC = TRUE))
setwd("C:/Personal_Cloud/OneDrive/Work/PublicCodesGit/Public/SpatialSurvivalOPSCR/")
save(myNimbleOutput, MCMCRuntime, file="samples.RData")
```

```
setwd("C:/Personal_Cloud/OneDrive/Work/PublicCodesGit/Public/SpatialSurvivalOPSCR/")
load("samples.RData")

#plot check 
chainsPlot(myNimbleOutput, var=c("N[1]","N[2]","N[3]","N[4]","N[5]"), line = N)
```

```
chainsPlot(myNimbleOutput, var=c("p0","sigma","mhW","mhH","tau","betaH","betaW"), 
           line= c(p0, sigma, mhW, mhH,tau, betaH, betaW))
```

#### REFERENCES

Bischof, Richard, Daniel Turek, Cyril Milleret, Torbjorn Ergon, Pierre Dupont, and Perry de Valpine. 2020. *nimbleSCR: Spatial Capture-Recapture (SCR) Methods Using ’Nimble’*.

de Valpine, Perry, Daniel Turek, Christopher J Paciorek, Clifford Anderson-Bergman, Duncan Temple Lang, and Rastislav Bodik. 2017. “Programming with Models: Writing Statistical Algorithms for General Model Structures with NIMBLE.” *J. Comput. Graph. Stat.* 26 (2): 403–13.

Dupont, P., C. Milleret, M. Tourani, H. Brøseth, and R. Bischof. 2021. “Integrating Dead Recoveries in Open-Population Spatial Capture–Recapture Models.” *Ecosphere* 12 (7): e03571. https://doi.org/https://doi.org/10.1002/ecs2.3571.

Milleret, Cyril, Pierre Dupont, Christophe Bonenfant, Henrik Brøseth, Øystein Flagstad, Chris Sutherland, and Richard Bischof. 2019. “A Local Evaluation of the Individual State-Space to Scale up Bayesian Spatial Capture–Recapture.” *Ecology and Evolution* 9 (1): 352–63. https://doi.org/10.1002/ece3.4751.

NIMBLE Development Team. 2020. “NIMBLE: MCMC, Particle Filtering, and Programmable Hierarchical Modeling.” https://doi.org/10.5281/zenodo.1211190.

Royle, J Andrew, and Robert M Dorazio. 2012. “Parameter-Expanded Data Augmentation for Bayesian Analysis of Capture–Recapture Models.” *Journal of Ornithology* 152 (2): 521–37.

Turek, Daniel, Cyril Milleret, Torbjørn Ergon, Henrik Brøseth, Pierre Dupont, Richard Bischof, and Perry de Valpine. 2021. “Efficient Estimation of Large-Scale Spatial Capture–Recapture Models.” *Ecosphere* 12 (2): e03385. https://doi.org/https://doi.org/10.1002/ecs2.3385.
