## Appendix 5 for "Estimating spatially variable and density-dependent survival using open-population spatial capture-recapture models"

Appendix 5: Using nimbleSCR to simulate from and fit Bayesian OPSCR models with spatial density dependent survival


### Appendix 5: Using nimbleSCR to simulate from and fit Bayesian OPSCR models with spatial density dependent survival

###### Cyril Milleret

#### 2022-02-25

In this vignette, we demonstrate how to use the nimbleSCR (Bischof et al. 2020) and NIMBLE packages (de Valpine et al. 2017; NIMBLE Development Team 2020) to simulate open-population spatial capture-recapture (OPSCR) data and fit flexible and efficient Bayesian OPSCR models with spatially explicit mortality. We assume that we have access to detections of individuals alive. The demographic model is parameterized using a hazard rate formulation and mortality is assumed to vary as a function of heterogeneous density. See details in Milleret et al…

```
## Load packages
library(nimble)
library(nimbleSCR)
library(basicMCMCplots)
library(raster)
library(coda)

## Functions will be uploaded into nimbleSCR
source("C:/Personal_Cloud/OneDrive/Work/PublicCodesGit/Public/SpatialSurvivalOPSCR/sampler_categorical_general1.R")
source("C:/Personal_Cloud/OneDrive/Work/PublicCodesGit/Public/SpatialSurvivalOPSCR/dcatState1Dead.R")
```

```
## Registering the following user-provided distributions: dcatState1Dead
```

```
source("C:/Personal_Cloud/OneDrive/Work/PublicCodesGit/Public/SpatialSurvivalOPSCR/calculateDensity.R")
```

#### 1. Simulate SCR data

##### 1.1 Habitat and trapping grid

As an example, we create a \(36 \times 36\) habitat grid with a resolution of 1 for each dimension. On the habitat, we center a \(28 \times 28\) trapping grid with also a resolution of 1 for each dimension.

```
r <- raster(nrows=9, ncols=9, xmn=0, xmx=36, ymn=0, ymx=36)
r[] <- 1
## Create habitat grid
coordsHabitatGridCenter <- coordinates(r)
colnames(coordsHabitatGridCenter) <- c("x","y")

## Create trapping grid
coordsObsCenter <- cbind(rep(seq(4.5, 31.5, by = 1), 28),
                         sort(rep(seq(31.5, 4.5, by = -1), 28)))
colnames(coordsObsCenter) <- c("x","y")

## Plot check
plot(r)
points(coordsHabitatGridCenter[,"y"] ~ coordsHabitatGridCenter[,"x"],
     pch = 1, cex = 1.5) 
points(coordsObsCenter[,"y"] ~ coordsObsCenter[,"x"], col="red", pch=16 ) 
par(xpd=TRUE)
legend(x = 5, y = 5,
       legend=c("Habitat window centers", "Trapping grid"),
       pt.cex = c(1.5,1),
       horiz = T,
       pch=c(1,16),
       col=c("black", "red"),
       bty = 'n')
```

```
habitatMask <- as.matrix(r)#matrix(1, nrow = 4, ncol= 4, byrow = TRUE)
```

##### 1.2 Rescale coordinates

To implement the local evaluation approach when fitting the SCR model (see Milleret et al. (2019) and Turek et al. (2021) for further details), we need to rescale the habitat and trapping grid coordinates so that each habitat cell is of dimension \(1 \times 1\). We also need to identify the lower and upper coordinates of each habitat cell using the ‘getLowAndUpCoords’ function.

```
## Rescale coordinates
scaledObjects <- scaleCoordsToHabitatGrid(
  coordsData = coordsObsCenter,
  coordsHabitatGridCenter = coordsHabitatGridCenter)

## Get lower and upper cell coordinates
lowerAndUpperCoords <- getWindowCoords(
  scaledHabGridCenter = scaledObjects$coordsHabitatGridCenterScaled,
  scaledObsGridCenter = scaledObjects$coordsDataScaled,
  plot.check = F)
```

We also set up the objects necessary to perform the local evaluation for the live detection using the ‘getLocalObjects’ function. Special care should be taken when chosing ‘dmax’ relative to \(\sigma\) (Milleret et al. 2019). Here we are using a value \(>3\*\sigma\) (see below for the \(\sigma\) chosen.

```
trapLocal <- getLocalObjects(habitatMask = habitatMask,
                             coords = scaledObjects$coordsDataScaled,
                             dmax = 1.8,
                             resizeFactor = 1,
                             plot.check = TRUE
)
```

##### 1.3 Define model code

Note that we do not provide ‘detNums’ and ‘detIndices’ arguments in the ‘dbinomLocal\_normal’ function as we wish to use this model to simulate data (see ‘?dbinomLocal\_normal’ for further details).

```
modelCode <- nimbleCode({
  ##--------------------------------------------------------------------------------------------
  ##-----------------------------## 
  ##------ SPATIAL PROCESS ------##  
  ##-----------------------------##
  tau ~ dgamma(0.001, 0.001)
  logHabIntensity[1:numHabWindows] <- mu[1:numHabWindows]
  sumHabInt <- log(sum(mu[1:numHabWindows]))
  ## FIRST YEAR 
  for(i in 1:M){
    s[i, 1:2,1] ~ dbernppAC(
      lowerCoords = lowerHabCoords[1:numHabWindows, 1:2],
      upperCoords = upperHabCoords[1:numHabWindows, 1:2],
      logIntensities = logHabIntensity[1:numHabWindows],
      logSumIntensity = sumHabInt,
      habitatGrid = habitatGrid[1:y.max,1:x.max],
      numGridRows =  y.max,
      numGridCols = x.max
    )
  }#i
  
  ## T>1 
  for(t in 2:n.years){
    for(i in 1:M){
      s[i, 1:2, t] ~ dbernppACmovement_normal(lowerCoords = lowerHabCoords[1:numHabWindows, 1:2]
                                                , upperCoords = upperHabCoords[1:numHabWindows, 1:2]
                                                , s = s[i, 1:2, t - 1]
                                                , sd = tau
                                                , baseIntensities = mu[1:numHabWindows]
                                                , habitatGrid =  habitatGrid[1:y.max,1:x.max]
                                                , numGridRows = y.max
                                                , numGridCols = x.max
                                                , numWindows= numHabWindows
      )
    }#i
  }#t
  
  ##--------------------------------------------------------------------------------------------
  ##-------------------------------## 
  ##----- DEMOGRAPHIC PROCESS -----## 
  ##-------------------------------##    
  ##INTERCEPT MORTALITY HR
  ## FIRST YEAR INCLUSION PARAMETER 
  gamma1 ~ dunif(0,1)
  omeg1[1:2] <- c(1-gamma1, gamma1)
  
  ## TIME SPECIFIC INCLUSION PARAMETER 
  for(t in 1:n.years1){
    gamma[t] ~ dunif(0,1)
  }#t
  
  

  betaPhi ~ dunif(-10,10)
  phi0 ~ dunif(-10,10)
  
  for(t in 1:n.years1){
   # for(r in 1:n.cells){
      phi[1:n.cells,t] <-  1-(exp(- exp(phi0 +  log(dens[1:n.cells, t]+1) * betaPhi)))
    #}
  }
  
  
  for(i in 1:M){ 
    z[i,1] ~ dcat(omeg1[1:2]) 
    for(t in 1:n.years1){
      z[i,t+1] ~ dcatState1Dead( z = z[i,t]
                                 , gamma = gamma[t]
                                 , phiSpatial = phi[1:numHabWindows,t]
                                 , s = s[i,1:2,t]
                                 , habitatGrid = habitatGrid[1:y.max,1:x.max]
      )
      
    }#i                                 
  }#t 
  
  ##---------------------------------------------------------------------------------------------   
  ##-----------------------------##
  ##----- DETECTION PROCESS -----## 
  ##-----------------------------##
  sigma ~ dunif(0,15)
  p0 ~ dunif(0,1)
  
  for(t in 1:n.years){
    for (i in 1:M){
      
      # ALIVE 
       y[i, 1:lengthYCombined,t] ~ dbinomLocal_normal(size = trials[1:n.traps],
                                                 p0 = p0,
                                                 s = s[i,1:2,t],
                                                 sigma = sigma,
                                                 trapCoords = trapCoords[1:n.traps,1:2],
                                                 localTrapsIndices = trapIndex[1:n.cells,1:maxNBDets],
                                                 localTrapsNum = nTraps[1:n.cells],
                                                 resizeFactor = resizeFactor,
                                                 habitatGrid = habitatIDDet[1:y.maxDet,1:x.maxDet],
                                                 indicator = isAlive[i,t],
                                                 lengthYCombined = lengthYCombined)
      
      
    }#i
  }#t
  
  ##---------------------------------------------------------------------------------------------         
  ##----------------------------------------## 
  ##---------- DERIVED PARAMETERS ----------##
  ##----------------------------------------##
  for(i in 1:M){ 
    isAlive[i,1] <- (z[i,1] == 2) 
    for(t in 1:n.years1){
      isAlive[i,t+1] <- (z[i,t+1] == 2) 
    }
  }
  
  for(t in 1:n.years){
    N[t] <- sum(isAlive[1:M,t])
  }#t
  
  # Density 
  for (t in 1:n.years) {
    dens[1:n.cells, t] <- calculateDensity(  s = s[1:M,1:2, t]
                                           , habitatGrid = habitatGrid[1:y.maxDet, 1:x.maxDet]
                                           , isAlive = isAlive[1:M,t]  
                                           , numWindows = n.cells
                                           , nIndividuals = M)#6
  }
  
})
```

##### 1.4 Define parameter values to simulate

```
# PARAMETERS
p0 <- 0.15 # We intentionally chose a large p0  to increase the size of the dataset and the convergence
sigma <- 2/res(r)[1] # Since we scaled the coordinates to the habitat, we also need to rescale the sigmaScaled 

M <- 650
n.individualsT1 <- 250
n.years <- 5
## ASSIGN SIMULATED VALUES 
phi0 <- 1.4

betaPhi <- -1
tau <- 1.5

recruitment <- 0.3  
## calculate the gamma 
gamma <- n.individualsT1/M
Recruit <- n.individualsT1*recruitment ## 40 % of recruitment 
NeverAlive <- n.individualsT1
  
for(t in 2:n.years){
    Navai  <- M- NeverAlive[t-1]
    gamma[t] <- Recruit/Navai 
    NeverAlive[t] <- NeverAlive[t-1] + Recruit
}
  
####
lengthYCombined <- 1 + trapLocal$numLocalIndicesMax*2
##
habCov <- as.numeric(scale(lowerAndUpperCoords$lowerHabCoords[,2])[,1])
habCovImage <- lowerAndUpperCoords$habitatGrid
habCovImage[] <- habCov[lowerAndUpperCoords$habitatGrid]
#check the habitat covariate. 
image(habCovImage)
```

```
### check survival probability relationship with density
dens <- log(c(1:10))
phi <- 1-(exp(- exp(phi0 +  dens * betaPhi)))
plot(phi ~ dens, ylim=c(0,1), type="b")
```

##### 1.5 Create data, constants and inits objects

```
nimConstants <- list(M = M,
                     n.years = n.years,
                     n.years1 = n.years-1,
                     n.traps = dim(scaledObjects$coordsDataScaled)[1],
                     y.max = dim(habitatMask)[1],
                     x.max = dim(habitatMask)[2],
                     y.maxDet = dim(trapLocal$habitatGrid)[1],
                     x.maxDet = dim(trapLocal$habitatGrid)[2],
                     resizeFactor = trapLocal$resizeFactor,
                     n.cells = dim(trapLocal$localIndices)[1],
                     maxNBDets = trapLocal$numLocalIndicesMax,
                     trapIndex = trapLocal$localIndices,
                     nTraps = trapLocal$numLocalIndices,
                     habitatIDDet = trapLocal$habitatGrid,
                     lengthYCombined = lengthYCombined,
                     numHabWindows = dim(lowerAndUpperCoords$lowerHabCoords)[1])


nimData <- list(trapCoords = scaledObjects$coordsDataScaled,
                trials = rep(1, dim(scaledObjects$coordsDataScaled)[1]),
                lowerHabCoords = lowerAndUpperCoords$lowerHabCoords,
                upperHabCoords = lowerAndUpperCoords$upperHabCoords,
                habitatGrid = lowerAndUpperCoords$habitatGrid,
                mu = rep(1, dim(lowerAndUpperCoords$lowerHabCoords)[1])
                )

# We set the parameter values as inits
nimInits <- list(sigma = sigma,
                 p0=p0,
                 tau=tau,
                 gamma = gamma[2:n.years],
                 gamma1 = gamma[1],
                 betaPhi = betaPhi,
                 phi0 = phi0)
```

##### 1.6 Create NIMBLE model

```
model <- nimbleModel( code = modelCode,
                      constants = nimConstants,
                      data = nimData,
                      inits = nimInits,
                      check = F,       
                      calculate = F)
```

##### 1.7 Simualte SCR data from the NIMBLE model

We first need to obtain the list of nodes that will be simulated. We used the ‘getDependencies’ function from NIMBLE. Using the ‘simulate’ function from NIMBLE, we will then simulate the activity center (AC) locations (‘s’), the state of the individual (‘z’) and SCR observation data (‘y’) given the values we provided for ‘p0,’ ‘sigma’ and ‘psi.’

```
# FIRST WE GET THE NODES TO SIMULATE
#nodesToSim <- model$getDependencies(c("s", "z"), self=T)
nodesToSim <- model$getDependencies(c("p0","sigma","phi0", "betaPhi","mu",
                                      "gamma","tau","gamma1"),
                                      self = F,
                                      downstream = T,
                                      returnScalarComponents = TRUE)
# THEN WE SIMULATE THOSE NODES 
set.seed(100)
model$simulate(nodesToSim, includeData = FALSE)
```

After running ‘simulate,’ the simulated data are stored in the ‘model’ object. For example, we can access the simulated ‘z’ and check how many individuals were considered alive and recovered at each time step :

```
N <- apply(model$z,2,function(x)sum(x==2))
N
```

```
## [1] 262 230 210 217 205
```

```
N.recoveredDead <- apply(model$z,2,function(x)sum(x==3))
N.recoveredDead
```

```
## [1]   0 114 202 262 351
```

```
# check average density 
mean(log(model$dens+1))
```

```
## [1] 1.210626
```

#### 2. RUN MCMC WITH NIMBLE

Here, we build the NIMBLE model again using the simulated ‘y’ as data. For simplicity, we used the simulated ‘z’ as initial values. Then we can fit the SCR model with the simulated ‘y’ data set.

```
myZ <- model$z
z <- zInits <- model$z
whichDet <- apply(model$y, 3, function(x) x[,1]>0 )
whichNotDet <- apply(model$y, 3, function(x) x[,1]==0 )

z[whichNotDet] <- NA
zInits[whichDet] <- NA

nimData$y <- model$y

nimData$z <- z
nimInits$z <- zInits
nimInits$s <- model$s


# CREATE AND COMPILE THE NIMBLE MODEL
model <- nimbleModel( code = modelCode,
                      constants = nimConstants,
                      data = nimData,
                      inits = nimInits,
                      check = F,
                      calculate = F)
model$calculate()
```

```
## [1] -22673.57
```

```
cmodel <- compileNimble(model)
cmodel$calculate()
MCMCconf <- configureMCMC(model = model,
                          monitors = c("p0","sigma","betaPhi","phi0", "N","dens",
                                         "gamma","tau"),
                          control = list(reflective = TRUE),
                          thin = 1)

###FIND WHICH Z NODES NEED TO BE REPLACED BY THE NEW SAMPLER. 
samplerConfList <- unlist(lapply(MCMCconf$getSamplers(),function(x) x$target))
zNodes <- samplerConfList[grep("z",samplerConfList)]
#find z nodes from the first year
zNodes <- zNodes[-grep( ", 1]",zNodes)]

MCMCconf
#remove samplers
MCMCconf$removeSamplers(zNodes)
for(i in 1:length(zNodes)){
  MCMCconf$addSampler(target = zNodes[i], type = 'sampler_categorical_general1',
                                  control=list("numCategories"= 3))
}

MCMC <- buildMCMC(MCMCconf)
cMCMC <- compileNimble(MCMC, project = model, resetFunctions = TRUE)
# RUN THE MCMC 
MCMCRuntime <- system.time(myNimbleOutput <- runMCMC( mcmc = cMCMC,
                                                             nburnin = 500,
                                                             niter = 5000,# need much longer run time (60000)
                                                             nchains = 2,
                                                             samplesAsCodaMCMC = TRUE))
```

#### 3. PLOT POSTERIORS

```
#plot MCMC 
chainsPlot(myNimbleOutput, var=c("N[1]","N[2]","N[3]","N[4]","N[5]"), line=N)
```

```
chainsPlot(myNimbleOutput, var=c("p0","sigma","betaPhi","phi0","tau"), line=c(p0, sigma, betaPhi, phi0, tau))
```

#### 3. IMPROVE MIXING BY CENTERING DENSITY

In a density-dependent survival OPSCR model, the log-density is used as a covariate for determining location- and time-specific survival rates. As expected for such model structures, use of this covariate induces a strong negative correlation between the posterior samples of the intercept and slope of the linear predictor (\(phi\_0\) and \(\beta\_{\phi}\), respectively). This particular model is complicated further by the fact that density itself is a non-constant latent variable, making it more difficult to address this posterior correlation and resulting in slow mixing for the density-dependent model. Here, we demonstrate that by centering the density (‘dens’), that is, try to make log(density+1) have a mean of 0, we can reduce the correlation between ‘betaPhi’ and ‘phi0’ and improve the mixing. We can’t do this exactly, because density is time- and location-dependent, but we can strive for 0-mean on average. From a preliminary run (in our case the one above), we can note the average of log(density+1) over time and space. For the specific dataset simulated in this vignette, mean(log(density)+1) = 1.2.

Now we can modify the model to center density such as:

```
for(t in 1:n.years1){
  phi[1:n.cells,t] <-  1- (exp(- exp(phi0 + (log(dens[1:n.cells, t]+1) - dens_offset) * betaPhi)))
}
```

Where the variable “dens\_offset” can be provided as a *constant value* in the nimConstants list.

```
mean(log(myNimbleOutput[[2]][,grep("dens", colnames(myNimbleOutput[[1]]))]+1))
```

```
## [1] 1.20669
```

```
nimConstants$dens_offset <- 1.2
```

Now we can rewrite the model code to center the density (‘dens’).

```
modelCode1 <- nimbleCode({
  ##--------------------------------------------------------------------------------------------
  ##-----------------------------## 
  ##------ SPATIAL PROCESS ------##  
  ##-----------------------------##
  tau ~ dgamma(0.001, 0.001)
  logHabIntensity[1:numHabWindows] <- mu[1:numHabWindows]
  sumHabInt <- log(sum(mu[1:numHabWindows]))
  ## FIRST YEAR 
  for(i in 1:M){
    s[i, 1:2,1] ~ dbernppAC(
      lowerCoords = lowerHabCoords[1:numHabWindows, 1:2],
      upperCoords = upperHabCoords[1:numHabWindows, 1:2],
      logIntensities = logHabIntensity[1:numHabWindows],
      logSumIntensity = sumHabInt,
      habitatGrid = habitatGrid[1:y.max,1:x.max],
      numGridRows =  y.max,
      numGridCols = x.max
    )
  }#i
  
  ## T>1 
  for(t in 2:n.years){
    for(i in 1:M){
      s[i, 1:2, t] ~ dbernppACmovement_normal(lowerCoords = lowerHabCoords[1:numHabWindows, 1:2]
                                                , upperCoords = upperHabCoords[1:numHabWindows, 1:2]
                                                , s = s[i, 1:2, t - 1]
                                                , sd = tau
                                                , baseIntensities = mu[1:numHabWindows]
                                                , habitatGrid =  habitatGrid[1:y.max,1:x.max]
                                                , numGridRows = y.max
                                                , numGridCols = x.max
                                                , numWindows= numHabWindows
      )
    }#i
  }#t
  
  

  
  ##--------------------------------------------------------------------------------------------
  ##-------------------------------## 
  ##----- DEMOGRAPHIC PROCESS -----## 
  ##-------------------------------##    
  ##INTERCEPT MORTALITY HR
  ## FIRST YEAR INCLUSION PARAMETER 
  gamma1 ~ dunif(0,1)
  omeg1[1:2] <- c(1-gamma1, gamma1)
  
  ## TIME SPECIFIC INCLUSION PARAMETER 
  for(t in 1:n.years1){
    gamma[t] ~ dunif(0,1)
  }#t
  
  

  betaPhi ~ dunif(-10,10)
  phi0 ~ dunif(-10,10)
  
  for(t in 1:n.years1){
   # for(r in 1:n.cells){
     phi[1:n.cells,t] <-  1- (exp(- exp(phi0 + (log(dens[1:n.cells, t]+1) - dens_offset) * betaPhi)))
    #}
  }
  
  
  for(i in 1:M){ 
    z[i,1] ~ dcat(omeg1[1:2]) 
    for(t in 1:n.years1){
      z[i,t+1] ~ dcatState1Dead( z = z[i,t]
                                 , gamma = gamma[t]
                                 , phiSpatial = phi[1:numHabWindows,t]
                                 , s = s[i,1:2,t]
                                 , habitatGrid = habitatGrid[1:y.max,1:x.max]
      )
      
    }#i                                 
  }#t 
  
  ##---------------------------------------------------------------------------------------------   
  ##-----------------------------##
  ##----- DETECTION PROCESS -----## 
  ##-----------------------------##
  sigma ~ dunif(0,15)
  p0 ~ dunif(0,1)
  
  for(t in 1:n.years){
    for (i in 1:M){ 
      # ALIVE 
       y[i, 1:lengthYCombined,t] ~ dbinomLocal_normal(size = trials[1:n.traps],
                                                 p0 = p0,
                                                 s = s[i,1:2,t],
                                                 sigma = sigma,
                                                 trapCoords = trapCoords[1:n.traps,1:2],
                                                 localTrapsIndices = trapIndex[1:n.cells,1:maxNBDets],
                                                 localTrapsNum = nTraps[1:n.cells],
                                                 resizeFactor = resizeFactor,
                                                 habitatGrid = habitatIDDet[1:y.maxDet,1:x.maxDet],
                                                 indicator = isAlive[i,t],
                                                 lengthYCombined = lengthYCombined)
      
      
    }#i
  }#t
  
  ##---------------------------------------------------------------------------------------------                                         
  ##----------------------------------------## 
  ##---------- DERIVED PARAMETERS ----------##
  ##----------------------------------------##
  for(i in 1:M){ 
    isAlive[i,1] <- (z[i,1] == 2) 
    for(t in 1:n.years1){
      isAlive[i,t+1] <- (z[i,t+1] == 2) 
    }
  }
  
  for(t in 1:n.years){
    N[t] <- sum(isAlive[1:M,t])
  }#t
  
  # Density 
  for (t in 1:n.years) {
    dens[1:n.cells, t] <- calculateDensity(  s = s[1:M,1:2, t]
                                           , habitatGrid = habitatGrid[1:y.maxDet, 1:x.maxDet]
                                           , isAlive = isAlive[1:M,t]  
                                           , numWindows = n.cells
                                           , nIndividuals = M)#6
  }
  
})
```

```
nimInits$phi0 <- -3 ## we modify the initial value.

model1 <- nimbleModel( code = modelCode1,
                      constants = nimConstants,
                      data = nimData,
                      inits = nimInits,
                      check = F,
                      calculate = F)
model1$calculate()
cmodel1 <- compileNimble(model1)
cmodel1$calculate()
MCMCconf1 <- configureMCMC(model = model1,
                          monitors = c("p0","sigma","betaPhi","phi0", "N","dens",
                                         "gamma","tau"),
                          control = list(reflective = TRUE),
                          thin = 1)

###FIND WHICH Z NODES NEED TO BE REPLACED BY THE NEW SAMPLER. 
samplerConfList <- unlist(lapply(MCMCconf1$getSamplers(),function(x) x$target))
zNodes <- samplerConfList[grep("z",samplerConfList)]
#find z nodes from the first year
zNodes <- zNodes[-grep( ", 1]",zNodes)]

MCMCconf1
#remove samplers
MCMCconf1$removeSamplers(zNodes)
for(i in 1:length(zNodes)){
  MCMCconf1$addSampler(target = zNodes[i], type = 'sampler_categorical_general1',
                                  control=list("numCategories"= 3))
}

MCMC1 <- buildMCMC(MCMCconf1)
cMCMC1 <- compileNimble(MCMC1, project = model1, resetFunctions = TRUE)
# RUN THE MCMC 
MCMCRuntime1 <- system.time(myNimbleOutput1 <- runMCMC( mcmc = cMCMC1,
                                                             nburnin = 500,
                                                             niter = 5000,# need much longer run time (60000)
                                                             nchains = 2,
                                                             samplesAsCodaMCMC = TRUE))
```

#### 3. PLOT POSTERIORS

```
#plot MCMC 
chainsPlot(myNimbleOutput1, var=c("N[1]","N[2]","N[3]","N[4]","N[5]"), line=N)
```

```
chainsPlot(myNimbleOutput1, var=c("p0","sigma","betaPhi","phi0","tau"), line=c(p0, sigma,betaPhi,phi0, tau))
```

We can compare the effective sample size of the the model without and without the density.

```
effectiveSize(myNimbleOutput1)[c("phi0","betaPhi")]
```

```
##     phi0  betaPhi 
## 78.16542 75.08951
```

```
effectiveSize(myNimbleOutput)[c("phi0","betaPhi")]
```

```
##     phi0  betaPhi 
## 11.97863 12.22744
```

Correlation between phi0 and betaPhi also decreased.

```
cor(myNimbleOutput1[[1]][,c("betaPhi")], myNimbleOutput1[[1]][,c("phi0")])
```

```
## [1] -0.9372544
```

```
cor(myNimbleOutput[[1]][,c("betaPhi")], myNimbleOutput[[1]][,c("phi0")])
```

```
## [1] -0.9937947
```

#### REFERENCES

Bischof, Richard, Daniel Turek, Cyril Milleret, Torbjorn Ergon, Pierre Dupont, and Perry de Valpine. 2020. *nimbleSCR: Spatial Capture-Recapture (SCR) Methods Using ’Nimble’*.

de Valpine, Perry, Daniel Turek, Christopher J Paciorek, Clifford Anderson-Bergman, Duncan Temple Lang, and Rastislav Bodik. 2017. “Programming with Models: Writing Statistical Algorithms for General Model Structures with NIMBLE.” *J. Comput. Graph. Stat.* 26 (2): 403–13.

Milleret, Cyril, Pierre Dupont, Christophe Bonenfant, Henrik Brøseth, Øystein Flagstad, Chris Sutherland, and Richard Bischof. 2019. “A Local Evaluation of the Individual State-Space to Scale up Bayesian Spatial Capture–Recapture.” *Ecology and Evolution* 9 (1): 352–63. https://doi.org/10.1002/ece3.4751.

NIMBLE Development Team. 2020. “NIMBLE: MCMC, Particle Filtering, and Programmable Hierarchical Modeling.” https://doi.org/10.5281/zenodo.1211190.

Turek, Daniel, Cyril Milleret, Torbjørn Ergon, Henrik Brøseth, Pierre Dupont, Richard Bischof, and Perry de Valpine. 2021. “Efficient Estimation of Large-Scale Spatial Capture–Recapture Models.” *Ecosphere* 12 (2): e03385. https://doi.org/https://doi.org/10.1002/ecs2.3385.
